# Do protein representations encode variant function? Diagnosing residue-level protein attributions via embedding-space gradients

**DOI:** 10.64898/2026.10.21.760763

**Authors:** Rana Barghout, Shengqing Wu, William Chau, Jose Manuel Barraza Chavez, Paola Driza, Radhakrishnan Mahadevan, Benjamin Sanchez-Lengeling

## Abstract

A common workflow in protein engineering involves building fitness predictors on protein language models (pLMs) to estimate a variant’s fitness and interpret which residues drive it. Attribution methods such as integrated gradients address this second goal by assigning residues contributions to the variant’s predicted fitness. However, without ground truth for residue importance, it is unclear whether these attributions reflect what the predictor learned about function. Here, we introduce **IGLOO**, a framework that scores residue-level attributions against synthetic fitness landscapes with known per-residue importance. Across experimental axes, high predictive accuracy did not guarantee faithful attributions. We identify model interventions that more than double attribution skill scores and raise TOPSIS by **29%**, with similar patterns holding in real ***α***-amylase and ***β***-lactamase datasets. IGLOO provides a protocol for probing what pLMs learn and a guide for choosing defaults when using residue attributions to pick sequence edits and build more trustworthy protein models.

## 1 Introduction

Protein language models (pLMs), trained on a large amount of natural sequences, learn residue-level representations that transfer to many prediction tasks. A common process for variant-effect prediction uses a pLM to embed each variant and trains a predictor on measured fitness labels [1, 2]. These predictors have shown strong performance on benchmarks such as ProteinGym [3], FLIP [4], and FLIP2 [5]. For protein engineering, ranking variants by predicted fitness is only part of the problem: practitioners also need to know *which residues drive function*, especially for enzymes whose structure or mechanism is uncharacterized; how are the positions for the next round of mutations chosen? Residue-level attribution is a common route to this kind of interpretation: converts a prediction into a per-residue score that can be compared with structure and tested by mutagenesis (Fig 1).

**Fig. 1.**
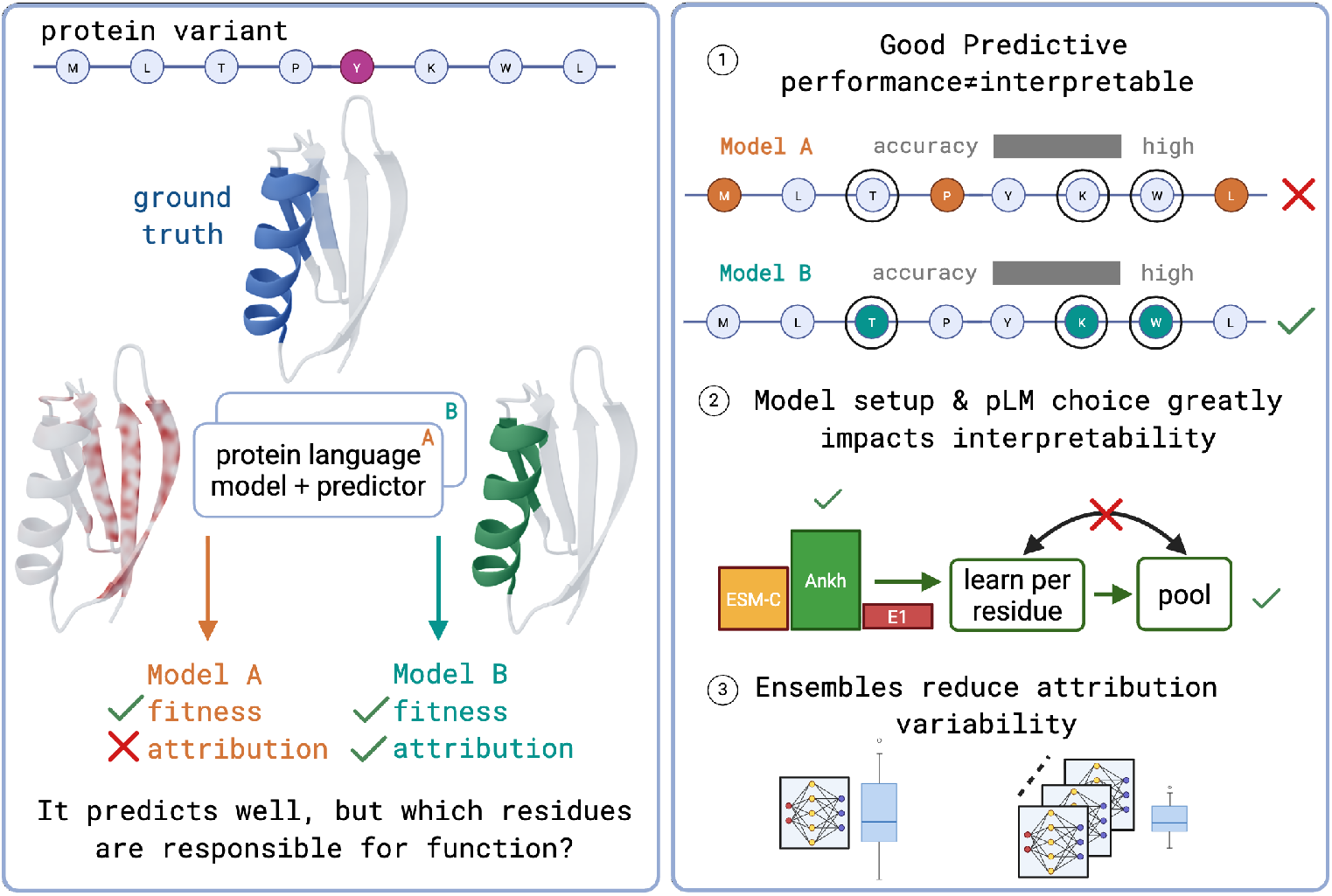
Good pLM-based fitness predictors can produce poor interpretability. (*Left*) pLM-based fitness predictors have been shown to have strong performance in learning fitness scores of different protein variants from their input protein sequence, but not all models that predict well are necessarily learning the right residue-level information related to that protein’s function. (*Right*) We find that (1) strong model predictive performance does not necessarily imply good interpretability; (2) model setup (pLM choice and layer as well as architecture and pooling layers) plays an important role in interpretability performance, even when predictive performance is comparable; (3) applying model ensembles can reduce the variability of interpretability performance.

Integrated gradients (IG) calculates these contributions by integrating gradients along a path between a reference input and the input being explained [6]. Applied to protein embeddings, IG produces an attribution score for each residue, allowing us to identify which parts of a protein sequence contribute to a model’s prediction. Reading such a map as an interpretation assumes that it reflects what the model learned and that the highlighted residues matter for the protein’s function. IG’s sensitivity and implementation invariance axioms provide a mathematical basis for attribution, but these qualities do not determine if the resulting maps identify biologically important residues. The scores can also depend on the reference input and integration path, and on how gradients behave in regions where the model output changes slightly [7–10].

Attribution methods have been applied to biological sequences to investigate features associated with predicted mRNA abundance [11], RNA splicing [12] and RNA protein-coding potential [13]. In proteins, interpretability studies have examined what pLM representations encode and, separately, which residues drive a trained model’s prediction [14]. For the former, InterPLM identified features in ESM-2 representations associated with biological concepts, including binding sites and structural motifs [15]. For the latter, Fazel et al. [16] compared explanation methods for interaction-site prediction and pLM embeddings. Wenzel et al. [17] extended IG to inspect internal representations of transformers fine-tuned for protein function prediction. Their analysis found statistically significant agreement between attribution maps and sequence annotations, including active sites and trans-membrane regions. Together, these studies show that pLM representations encode biological information and that attribution methods can recover annotated regions for particular prediction tasks. A pLM fitness predictor contains both a pretrained representation and a trained head, so an embedding-space attribution could reflect either.

However, high predictive accuracy does not imply that these attribution maps capture biological information. Del Giudice [18] calls this the prediction–explanation fallacy: a model optimized to predict is often used to explain, even though models that predict equally well can imply conflicting explanations. A recent study of protein allergenicity found that pLM-based classifiers predicted well, yet their residue attributions were no better than random for annotated epitopes [19]. Furthermore, for pLM-based fitness predictors, the setting most used in protein engineering, no study has quantified, to our knowledge, whether residue attributions recover the residues that determine fitness, or which modeling choices decide whether they do.

Evaluating attributions therefore requires evidence beyond prediction accuracy. In computer vision and natural language processing, pixel-flipping and ROAR test how changing or removing highly scored features affects predictions [20, 21], while ERASER also compares explanations with human annotations [22]. For molecular graphs [23], benchmarks with known ground-truth attributions compare attribution methods across model architecture [24]. Similar evaluations in proteins require meaningful sequence changes and a definition of residue importance [25]. Deep mutational scanning measures variant effects, but assigning importance to each position requires deciding how to combine different substitutions and sequence contexts. Known functional sites provide another reference, although they may not capture every residue that affects the measured fitness.

We therefore ask under what conditions residue-level attributions from a pLM fitness predictor are trustworthy, and how practitioners can tell whether their own pipeline meets those conditions (Fig 1, left). We call an attribution *trustworthy*, meaning reliable enough to act on when choosing sequence edits, when it (i) carries signal above chance on a metric capable of detecting it; (ii) depends on both the protein embedding and the trained predictor; and (iii) agrees with known functional residues in real proteins.

Here we introduce **IGLOO** (**I**ntegrated **G**radients for **LO**cational sensitivity in pr**O**teins), a framework that measures how well residue-level IG attributions from pLM-based fitness predictors recover known functional residues. Our contributions are:

1. **A benchmark with known residue-level ground truth**. IGLOO makes attributions quantifiable by defining residue-level ground truth in synthetic fitness landscapes built on three proteins with distinct topologies (GB1, p53 and T4 lysozyme) under additive, epistatic, and nonlinear fitness models.
2. **A single attribution score**. It trains predictors on embeddings from three pLM families (ESM-C [26], Profluent E1 [27], and Ankh [28, 29]) and combines five attribution metrics into a single TOPSIS score.
3. **Model interventions that improve attribution quality and trustworthiness**. With IGLOO, we show that predictive accuracy does not guarantee trustworthy attributions (Fig 1, right), and we identify three interventions that make attributions more trustworthy:

- the choice of pLM;
- the setup and ordering of modeling layers;
- ensembling, which reduces variability.

We then apply IGLOO to two enzymes with experimental fitness data and characterized active sites, *α*-amylase and *β*-lactamase, to test criterion (iii) on real proteins. IGLOO is open and extensible to new pLMs, predictor heads and proteins. It offers the community a common standard for asking what pLM-based fitness predictors learn about protein function, and gives practitioners a concrete test of when residue-level attributions can and cannot be trusted.

## 2 Results

### 2.1 A synthetic benchmark makes protein attribution quality measurable

To evaluate residue-level attributions from embedding-space integrated gradients, we construct synthetic datasets based on three proteins: GB1, T4 lysozyme, and p53. For each protein, we assign literature-derived importance weights to individual residues and use these weights as ground truth for attribution evaluation (Fig 2, 1). We generate fitness labels using additive, epistatic, and nonlinear models, yielding nine protein–landscape combinations. For each combination, we vary label noise across nine levels, from clean to fully randomized labels, to examine how attribution quality changes with predictive performance.

**Fig. 2.**
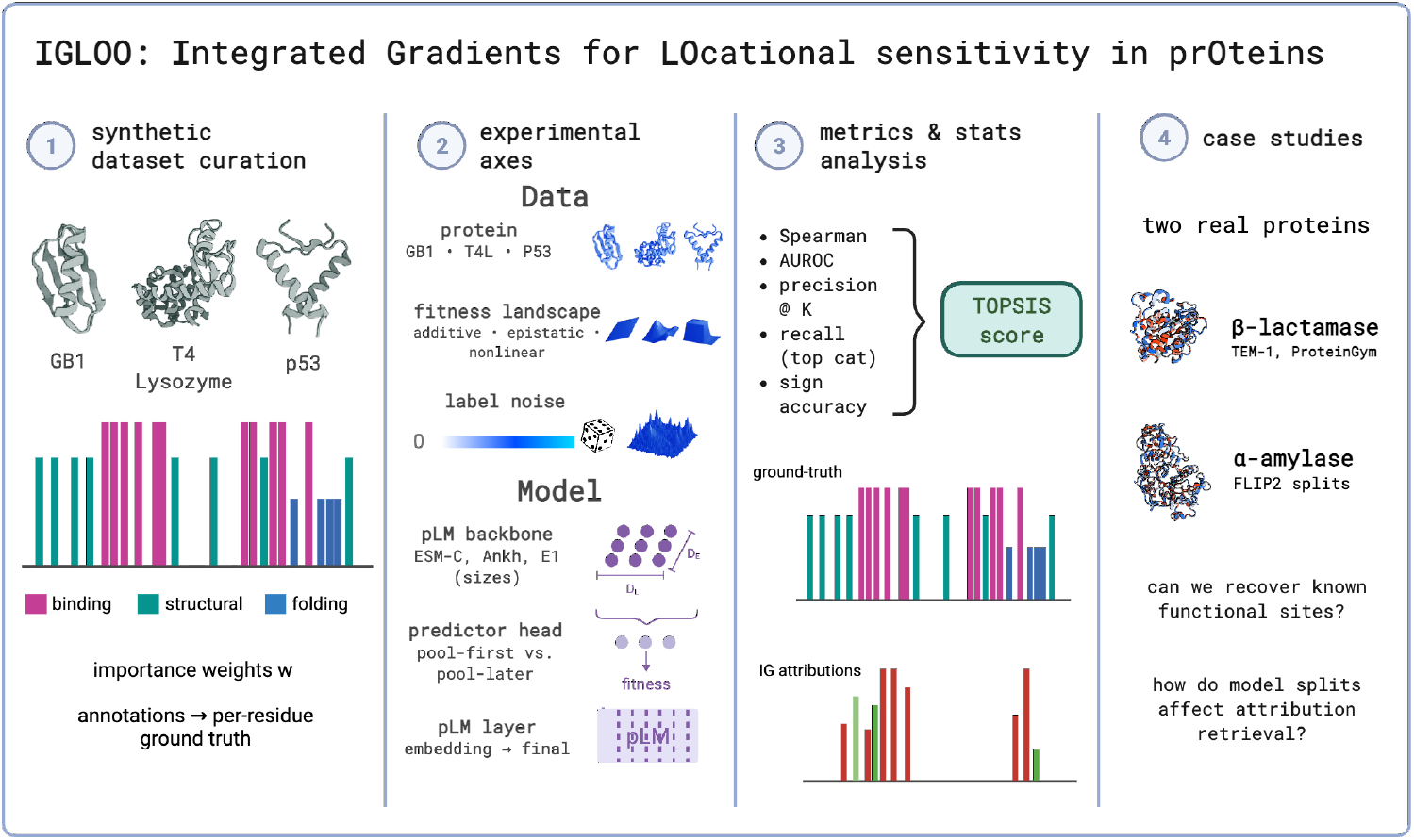
IGLOO experimental setup. (**1**) We curate synthetic datasets from three real proteins (GB1, T4 Lysozyme, and p53) by assigning importance weights to functionally annotated sites and use them as per-residue ground truth values for attribution comparison. (**2**) We test attribution performance across six different axes. In the data, we test the 3 different proteins across 3 different fitness landscapes and vary label noise to test attribution performance across a range of model performance. In the model, we test 7 different pLMs in 3 different pLM families, different predictor architecture options that shuffle pooling location, and also different pLM representation layers. (**3)** We aggregate 5 different evaluation metrics into one TOPSIS score to evaluate our obtained IG attributions against ground-truth weights. (**4**) Two real protein datasets are used as case studies for our pipeline to demonstrate real protein design workflows for IGLOO.

We compare attribution quality across three pLM families (ESM-C, Ankh, and E1) and their different model sizes. We also compare pool-first predictors, which apply attention pooling directly to the pLM embeddings, with pool-later predictors, which learn per-residue representations before pooling. We further evaluate intermediate pLM layers as sources of protein representations for integrated gradients (Fig 2, 2).

We evaluate the resulting attributions against the ground-truth residue weights using five metrics (Section 4.6). We combine these metrics into a single TOPSIS score to compare configurations across the evaluation criteria (Fig 2, 3). We then apply IGLOO to two real protein case studies (Fig 2, 4), discussed in Section 2.3.

**Fig. 3.**
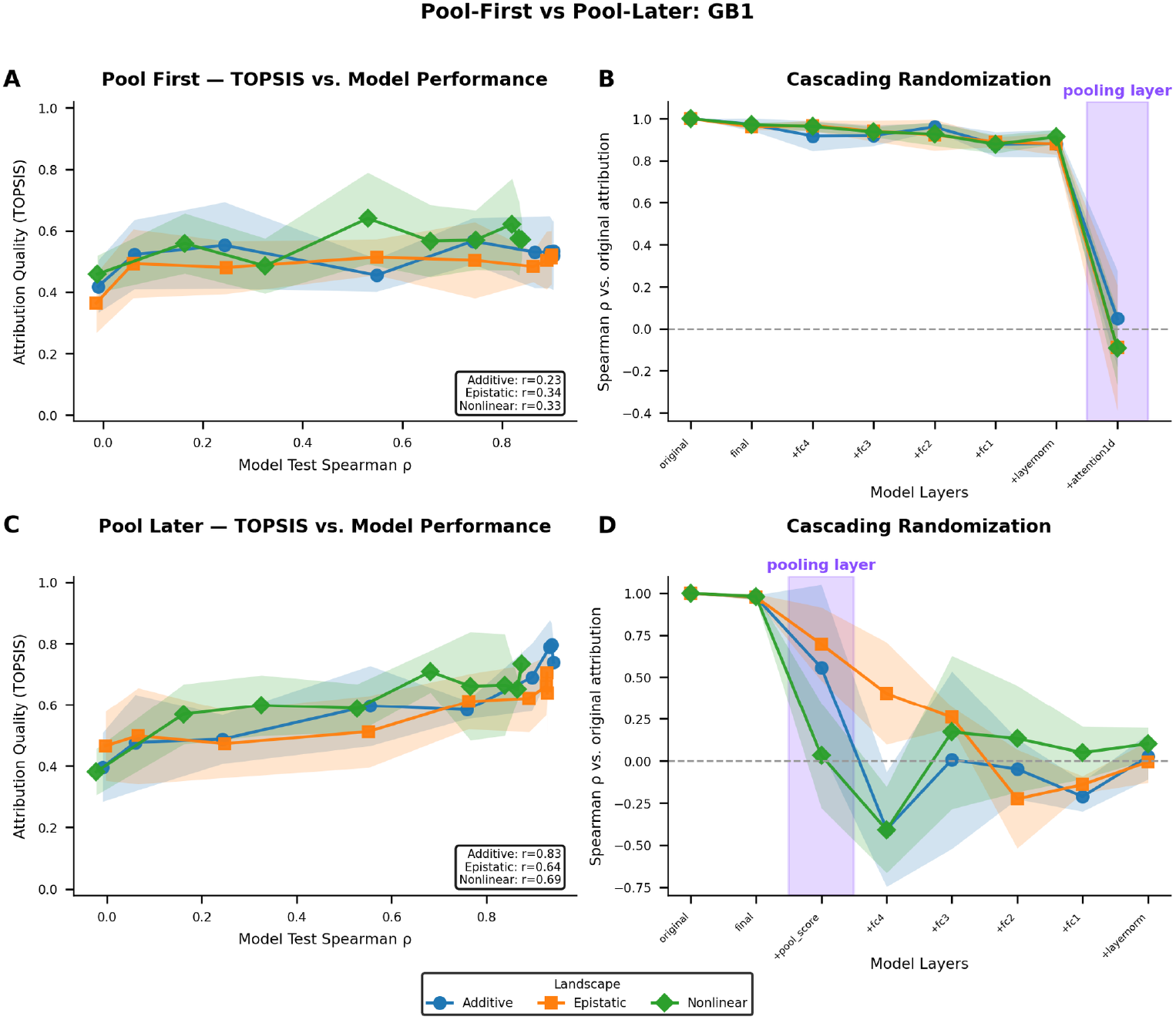
Correct model intervention improves model interpretability. (**A**) Attribution quality TOPSIS score vs. model test Spearman performance. All landscapes show weak correlation between interpretability and model predictive performance. (**B**) We apply incremental per-layer weight randomization (cascading randomization) on the predictor architecture in a top-to-bottom manner and find that the only layer that has a large effect on attribution (y-axis shows the Spearman correlation between the original attribution and attribution after layer weight randomization) is the attention pooling layer applied on the pLM embedding (**C)** TOPSIS score vs model test Spearman performance correlation improves when we apply this pool-later architecture intervention. (**D**) When we apply pooling later, we see that intermediate layers have a cumulative effect on attribution.

**Fig. 4.**
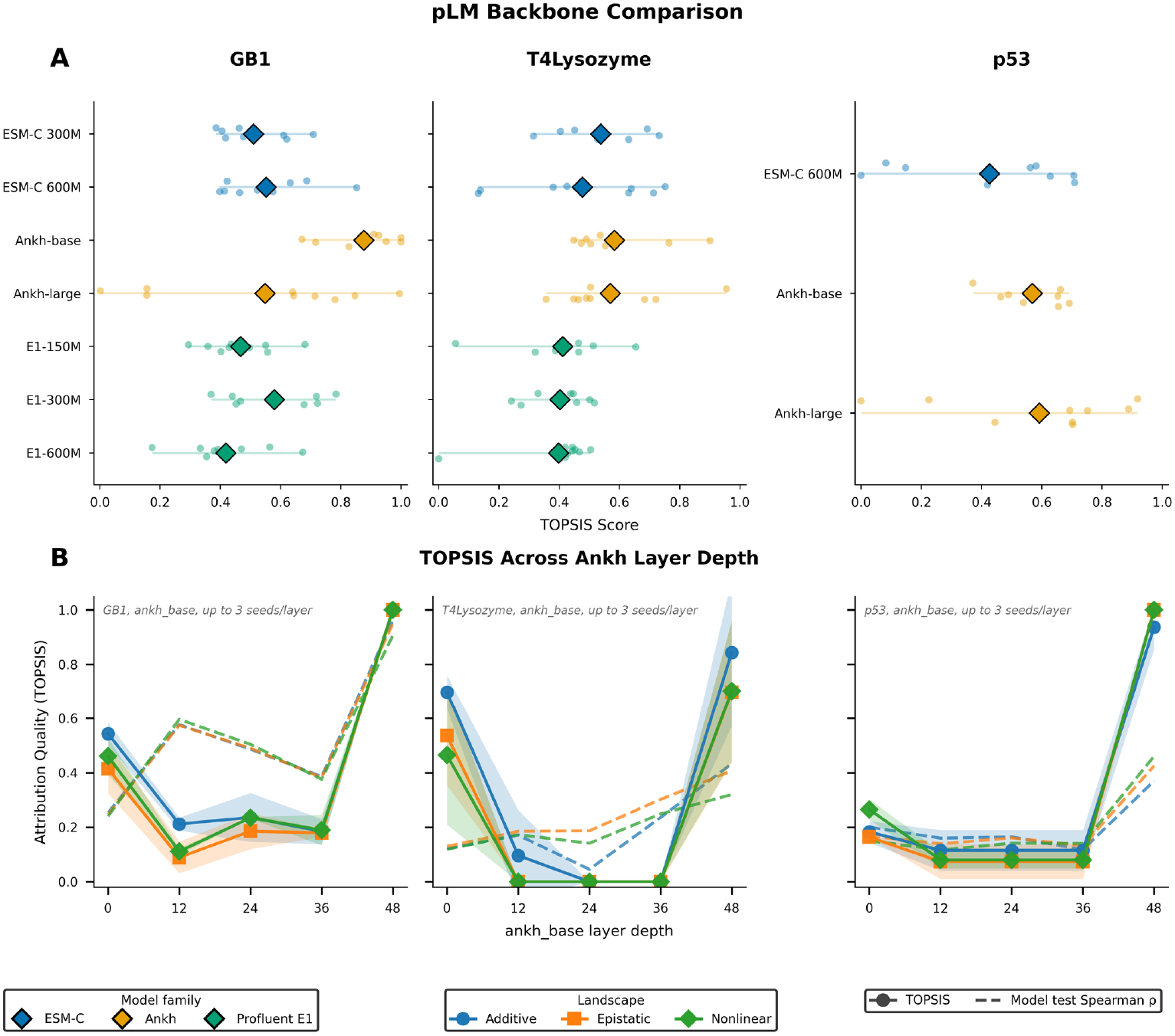
pLM backbone choice affects attribution quality. (**A**) Across our tested proteins, we see that the Ankh family of models produce the best performing attributions when aggregated across landscapes. *Only the top three pLMs from GB1 & T4 Lysozyme were tested for the p53 datasets to reduce computational cost. (**B**) Ankh-base intermediate layer representations don’t provide better attributions; the attribution performance collapses in these intermediate pLM representations, even though model test performance fluctuates.

### 2.2 Experiments in a controlled setting

#### 2.2.1 Attribution quality weakly tracks accuracy in standard attention-pooled predictor

We train a fitness predictor on ESM-C 300M residue embeddings, using attention pooling to compress the length dimension and compile into a single, dimension-invariant representation for each variant. This approach pools residue embeddings then sends the pooled variant representations directly to the fitness predictor MLP, and was similarly done in FLIP, CLEAN, UniKP, and XAllergen [4, 19, 30, 31]. We vary the amount of noise added to the fitness labels to obtain models with a range of test performances. Across all three fitness landscapes, attribution quality shows little improvement as predictive performance increases, and does not noticeably exceed that of a model that has not learned the relationship between sequence and fitness (Fig 3 panel A).

#### 2.2.2 Attributions are determined by the pooling layer

In order to better understand why attribution performance was not improving as the predictor learned the fitness distributions of our variants, we ran a cascading randomization experiment, introduced by Adebayo et al. [32], on our fitness predictor (Fig 3 panel B). We observe that attribution rank correlation with our original attributions (prior to layer weight randomization) stays high (∼0.8–1.0) while every MLP layer and the layer norm are randomized, and collapses only when the attention-pooling layer is randomized.

#### 2.2.3 Moving learned computation before pooling strengthens link to attribution accuracy

Based on the behavior we observed in the cascading randomization experiment in Fig 3 panel B, we instead tested architectures that apply residual per-residue MLP before the attention pooling to test whether the IG can take advantage of these learnt gradients in retrieving attributions (Appendix B.1). We tested three alternative pool-later architectures (Residual Pre-MLP, Per-residue + Attn Pool (AttnPool), and Per-residue + Mean Pool (MeanPool)) as well as a one-hot encoding baseline (further described in Methods section 4.3 and Appendix B) and observed that per-residue processing before pooling gives stronger correlation between attribution quality and model predictive accuracy, with Pearson’s *r* going from 0.23 to 0.83 in the GB1 additive fitness landscape (Fig 3 panel C). We also observe that the cascading randomization has a cumulative degradation effect on the attribution compared to the original attribution obtained when we apply this pool-later intervention (Fig 3 panel D). Similar behaviors were observed in other proteins and landscapes; results demonstrating this attribution–model performance behavior for our three proteins across the landscapes are presented in Appendix D, Fig D.1. Full metric breakdown across the different architectures is presented in Fig D.2.

#### 2.2.4 pLM backbone strongly affects attribution quality, and larger models don’t necessarily help

Next, to test whether different pLM representations had a large influence on attribution quality, we applied IGLOO to our synthetic datasets across seven different pLMs in the ESM-C, Ankh, and E1 families paired with the pool-later predictor architecture. We observe that the Ankh model is the most consistent backbone across landscapes, with the smaller Ankh-base model outperforming the larger Ankh-large model on GB1 across landscapes (Fig 4 panel A), demonstrating that model scale does not necessarily improve attribution. The full backbone × pool-later architectures TOPSIS scores are presented in Appendix D, Figure D.3. The performance breakdown per metric for a strong pool-later architecture against baselines is presented in Appendix Figure D.4.

#### 2.2.5 Within a pLM, intermediate layers predict ‘well’ but attribute poorly

Motivated by the finding that intermediate pLM representations can sometimes lead to better function prediction performance in certain applications [33–36], we test whether IG applied to representations from intermediate layers in the Ankh-base pLM also demonstrate this behavior. We find that intermediate layers show a collapse in attribution quality in all protein datasets (Fig 4), and that the best attribution performance is obtained from the last layer representation in the model. Interestingly, we also observe that there is a disconnect between attribution performance and model predictive performance in all three proteins as well, where model predictive performance can reach Spearman’s *ρ* of 0.6 while attribution TOPSIS score collapses to *<* 0.3.

#### 2.2.6 Model ensembles reduce attribution variability

From previous experiments, we observed that the quality of model attribution can vary drastically (Fig 5). To measure this variability and how it shrinks when models are ensembled, we trained 10 independently seeded Ankh-base + per-residue attention pooling models on GB1. For each ensemble size *K* ∈ {1, …, 10}, we computed the standard deviation of the TOPSIS score two ways: (i) 300 bootstrap draws of *K* seeds with replacement from the pool of 10 trained seeds, and (ii) the exact standard deviation computed from every distinct group of *K* seeds, with no resampling. To confirm that the resulting decrease in standard deviation reflects genuine ensembling behavior rather than an artifact of how the ensembles were sampled, we checked that both approaches agree. They produce closely matching curves that track the theoretical 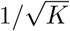 line (Fig 5, panel B), confirming that the ten trained seeds behave like independent sources of noise. The inter-quartile range of the TOPSIS score is also shifted with a 10-seed ensemble (Fig 5, panel C).

**Fig. 5.**
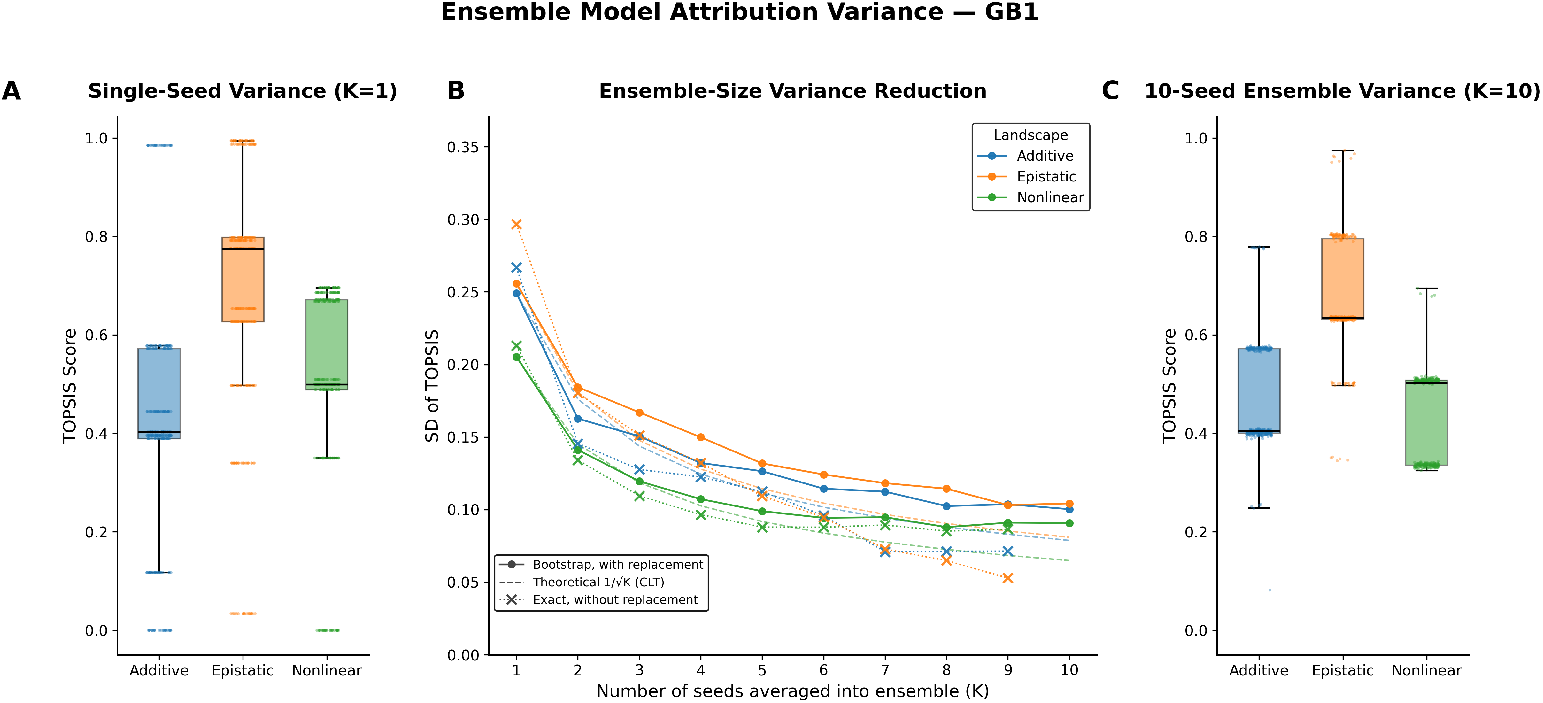
Ensemble models produce more stable attributions. (**A**) Box-plot of the TOPSIS score for a single-seed model across the different GB1 fitness landscapes. (**B**) Standard deviation of the TOPSIS score vs. number of models averaged into the ensemble (*K*). Solid line: 300 bootstrap draws of *K* seeds *with replacement* from the pool of 10 trained seeds. Dotted line: the exact standard deviation from exhaustively enumerating all 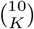 distinct size-*K* subsets of the 10 seeds *without replacement* (undefined, and omitted, at *K* = 10, where only one such subset exists). Dashed line: the theoretical 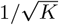 rate predicted by the Central Limit Theorem for the standard deviation of a sample mean of *K* independent draws.(**C**) Box-plot of the TOPSIS score for a 10-seed ensemble model across the different GB1 fitness landscapes. All data points were sampled from the Ankh-base with Per-residue + Attn Pool model.

### 2.3 Deploying interpretability in real settings

To explore how these observed patterns translate to real protein datasets, we applied IGLOO to two real protein case studies: an *α*-amylase site saturation mutagenesis dataset split four ways from FLIP2 [5], and four independent *β*-lactamase deep mutational scanning datasets from ProteinGym [3]. For the *α*-amylase datasets, the top twenty-five positions from our IG attributions were selected and compared against our per residue level labels using the Ankh-base + pool-later model. As observed in the synthetic datasets, we see that the split that yielded the highest model test performance did not yield the best attribution performance (Fig 6). For the `mutation by` mutation split (far left in Fig 6), the model’s held-out Spearman *ρ* is 0.66*±*0.01 and the recall values range from 0.45*±*0.05 to 0.56*±*0.01, while the `close to far` split has a model held-out Spearman *ρ* of 0.43*±*0.02 but shows the best recall for the evolutionarily conserved sites with attributions retrieving 0.67*±*0.01 of the protein’s sites. Full performance across different splits for the *α*-amylase dataset are presented in Appendix Table E.1. Pool-later models also outperformed pool-first models on attribution in every split, even where pool-first models predicted activity as well or better (Appendix Figure E.1). Next, we wanted to understand how IGLOO’s attributions would perform in an engineering setting, where the attributions were used to guide perturbations in the wild-type TEM-1 *β*-lactamase sequence and their effect was predicted using Boltz-2 [37] affinity and Rosetta cartesian ΔΔ*G* scores. We observe that across the 35 total substituted position and ligand combinations, 63% combinations led to the expected effect when compared against binding affinity and 69% of the combinations led to the expected effect when compared against stability (Appendix Tables E.2 and E.3), furthering interest in future exploration on how IGLOO can be used in the protein engineering design loop.

**Fig. 6.**
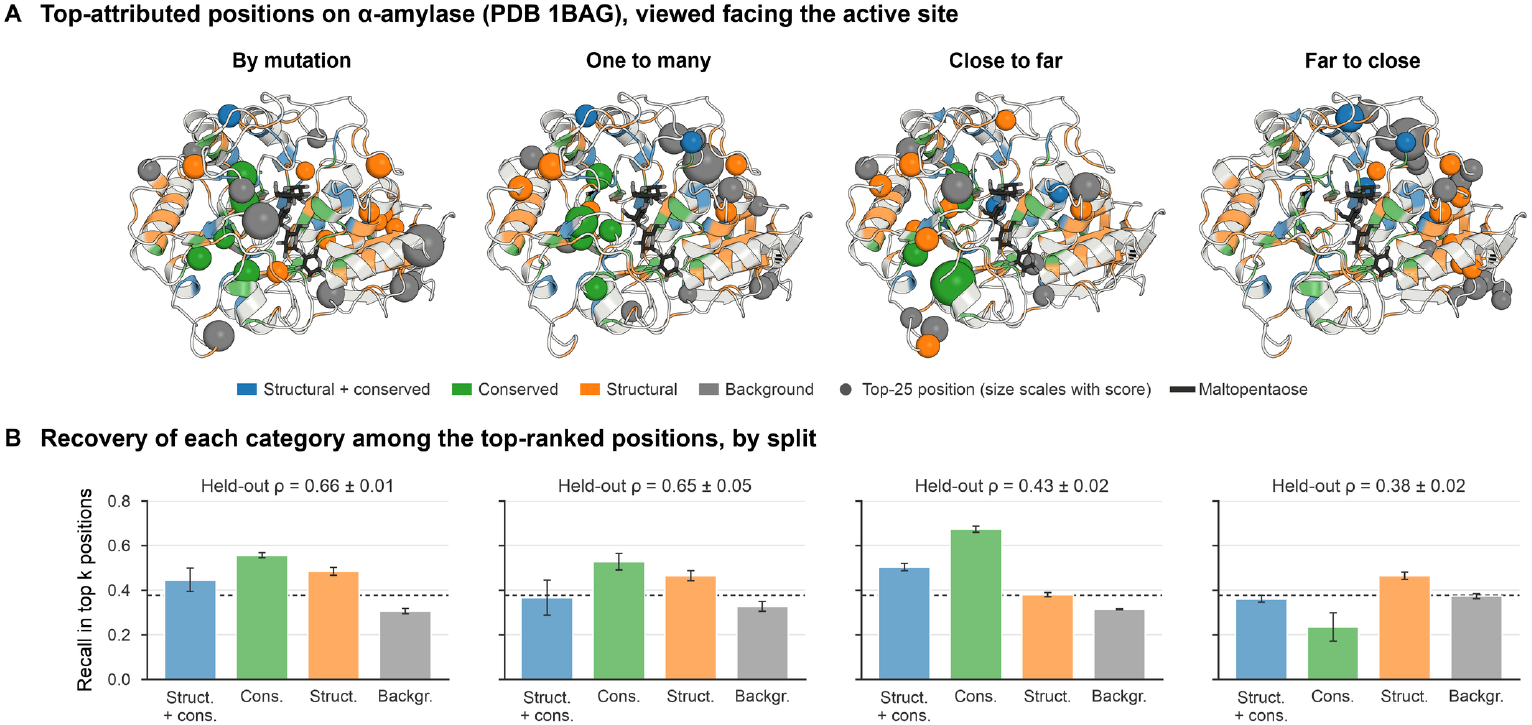
Different splits recover different functional categories in *α*-amylase. (**A**) Top-attributed positions mapped onto *α*-amylase (PDB 1BAG) for each FLIP2 split. The cartoon is colored by position category, defined in Section 4.9.4. Spheres mark the 25 positions with the highest ensemble attribution score (mean |*IG*|*×* reliability ratio across 5 seeds), colored by category and sized by score. (**B**) Fraction of each category’s positions ranked in the top *k* = 160 by attribution magnitude, where *k* is the number of annotated positions. Bars show the mean *±*s.d. over 5 seeds; the dashed line marks the random expectation (*k/N* =0.38). Held-out *ρ* is the Spearman correlation between predicted and measured activity on each split’s test set. Attribution metrics and TOPSIS scores are given in Appendix Table E.1.

## 3 Discussion

Here, we introduced IGLOO, a quantitative framework to benchmark and diagnose the fidelity of residue-level attributions in pLM-based fitness predictors. By evaluating seven pLM backbones across diverse synthetic and empirical fitness landscapes, our analysis reveals an important finding for the protein engineering community: high predictive accuracy (Spearman’s *ρ*) does not guarantee that a model relies on interpretable, biologically grounded residue-level features. This disconnection highlights the importance of going beyond the use of feature attributions for targeted sequence editing based solely on a model’s regression performance. Instead, IGLOO reveals that attribution quality is highly sensitive to deliberate architectural choices, specifically the location of the pooling layer and the choice of the embedding backbone. Our architectural sweep demonstrates that the arrangement of the prediction head dictates how a model distributes gradient information. When a predictor pools per-residue pLM embeddings (e.g. via a learned attention pool) before any predictor layers, the downstream head contributes only a single, shared gradient direction with respect to the pooled representation. Each position’s gradient is then this shared vector rescaled by position-specific scalars set by the shallow pooling layer, so per-residue attributions are largely determined by the pooling weights and the embedding displacements. The learned head can re-weight positions only along this one shared direction and cannot assign importance to individual positions independently. On the other hand, placing learned per-residue layers before pooling makes each position’s gradient a nonlinear function of its own (context-aware) pLM embedding, so attribution can be reallocated to individual positions. Tracing attributions through our framework yields a practical guide: model architecture has the largest effect on attribution quality, followed by backbone selection, with multi-seed ensemble models further reducing variance. Namely, we find that a ‘pool-later’ per-residue head with attention pooling operating on Ankh-base final-layer embeddings consistently yields the most reliable attributions, nearly doubling per-metric skill scores relative to the conventional ‘pool-first’ baseline (AUROC skill 0.42 → 0.88; Spearman skill 0.27 → 0.56, averaged across all three synthetic proteins) and raising composite faithfulness (TOPSIS) by ∼29% (0.51 → 0.65).

To explain why intermediate pLM layers and larger models often failed to improve attribution quality, we present two observations. First, attribution collapsed in the intermediate layers of Ankh-base (layers 12-36), and recovered fully at the final layer. Ankh’s intermediate hidden states are the unnormalized residual stream of a T5 encoder. Only the final layer passes through the encoder’s output normalization. The intermediate states are dominated by a few massive-activation dimensions (∼10^3^*×* the median dimension), which degraded either model training (T4 Lysozyme, p53) or the alignment of attributions with ground truth (GB1). The embedding layer (layer 0) is a context-free token lookup, equivalent to a linear projection of a one-hot encoding. Its attributions are therefore confined to mutated positions by construction, consistent with our one-hot encoding baseline. Normalizing intermediate layers before that head would test whether this dip is a scaling artifact. Second, attribution quality did not necessarily increase with parameter count, either across backbone families or between Ankh-base and Ankh-large, which share depth and differ only in width. This suggests that factors other than scale (possibly training data, model architecture, and pretraining setup) dictate how much residue-level signal can be extracted. These factors are confounded across the backbones we tested.

Based on these findings, we propose a diagnostic checklist for practitioners utilizing residue attributions to guide sequence edits:

- **Process before pooling:** Implement learned, per-residue layers prior to any pooling or global aggregation steps
- **Prioritize Ankh backbones:** Leverage Ankh embeddings as a robust default for feature extraction over ESM-C or E1 embeddings
- **Ensemble predictors:** Average attributions across an ensemble of 5 ™ 10 independent models; given the low computational cost of training downstream heads, this provides an accessible boost to attribution stability
- **Benchmark against absolute baselines:** Report attribution quality rigorously against absolute chance, rather than relying exclusively on relative rankings when evaluating your own attributions

Because attribution is cheap relative to training (IG for one variant costs ∼0.1 s on a single GPU verus ∼10 mins to train the downstream head once), we see no computational reason to omit attribution analysis whenever these predictors are deployed for sequence-editing decisions. Moreover, applying the ‘pool-later’ intervention does not increase model parameters compared to the ‘pool-first’ approach.

We used the *α*-amylase system as a case study to test our prior findings, and the *β*-lactamase system to demonstrate a potential use case of IGLOO. As both of the case studies’ definitions of fitness involve multiple complex intracellular processes, we simplified the contributing components of fitness to protein abundance and catalytic activity. This means that the attributions obtained from IG may be finding signals that are out of scope of our current methodology. Regardless, the results of the *α*-amylase match our finding that model prediction accuracy doesn’t guarantee faithful attribution. We also observe that the far to close split does not agree well with the other splits, with a notably low amount of conserved residues. This can possibly be credited to the fact that all residues within 7.3 Åof the catalytic site were withheld, and thus residues conserved for their effects on catalysis and binding were not seen by the model. This means that IGLOO cannot properly assign attributions to positions it has not seen. For the *β*-lactamase case study, taking the top 5 attributed positions of each dataset produced 18 unique positions. We observe that about two thirds of all tested mutants demonstrate an effect that matches the attribution sign. However, the mutants with effects that did not match the attribution sign do not necessarily suggest that the positions were misattributed, as the proxy measures of fitness do not cover all factors that contribute to the true definition of fitness. Interestingly, nine unique positions found have annotated functions, with two for catalysis, one for substrate binding, one that participates in disulfide bridge, and five for structural stability. Three positions found are directly next to positions with annotated functions, with two being next to catalytic residues and one next to a structural residue. There are two positions shared within the top 5 attributed positions of the datasets: Ala235 catalytic residue is shared between the Deng et al. and Stiffler et al. datasets, which test on ampicillin and both ampicillin and cefotaxime respectively, and the Val29 structural residue is shared between the Firnberg et al. and Jacquier et al. datasets, which test on ampicillin and amoxicillin respectively. This suggests that there is agreement of attribution signals even when the datasets have independent methodologies and slightly differing definitions of function.

We acknowledge several limitations in the current implementation of the framework. First, the synthetic ground truths generated within IGLOO rely on specific assumptions, the first being that BLOSUM substitution matrices adequately capture baseline substitution weights and that epistatic interactions can be entirely reconstructed from physical residue contacts (≥2 mutations). While our structural sweeps rigorously covered three distinct proteins (GB1, T4 Lysozyme, and p53) across nine protein–landscape combinations, this sample size bounds our global statistical power. Furthermore, the TOPSIS framework used for multi-criteria optimization remains inherently relative, and the residue edits nominated by our pipeline were validated *in silico* rather than through wet-lab experimental expressions. Lastly, across both real protein scenarios, we used predictive models (Rosetta cartesian ΔΔ*G* and Boltz-2) to compute the mutational effects on stability and binding affinity. These scores inform us of the effects on proteins, which maps to experimentally determined stability (ΔΔ*G*) and catalysis (binding affinity) with inherent errors coming from the modeling methods. Stability and catalysis are themselves a simplified representation of enzyme or organismal fitness, as there are many other non-protein factors in play that are not within scope. Nevertheless, our real protein scenarios demonstrate the potential of IGLOO when applied to proteins. Looking forward, we envision IGLOO serving as an open-access diagnostic standard for the community. As new, multibillion parameter pLMs and novel predictor architectures are continuously introduced, IGLOO can be immediately deployed to pre-screen whether these models are strong contenders for mechanistic interpretation and attribution-guided design. Expanding the framework to encompass a wider diversity of protein topologies and validating IGLOO-selected mutations through direct experimental assays will represent essential next steps toward fully automating trustworthy, loop-closing protein design.

## 4 Methods

### 4.1 Synthetic Data

To evaluate IG attribution quality against known ground truth, we construct synthetic fitness datasets for three proteins spanning a range of sequence lengths, annotation densities, and functional complexities: **GB1** (P06654: 56 amino acids (aa), IgG-binding B1 domain), **T4 Lysozyme** (P00720: 164 aa, bacteriophage lytic enzyme), and **p53** (P04637: 393 aa, multi-domain tumor suppressor). Each residue is assigned a functional category and importance weight *w*_*i*_ (active site, binding site, structural, folding, signaling, or metal binding, highlighted in Table A.1). We implement three fitness functions of varying complexity. All three landscapes are defined in terms of the set of mutations *M*(**x**) carried by variant **x** relative to the wild type (WT). Each mutation *k* ∈ *M*(**x**) substitutes WT residue *a*_*k*_ with residue *b*_*k*_ at position *i*_*k*_. Each mutation is assigned a substitution penalty derived from BLOSUM62,

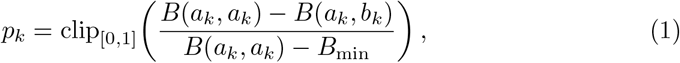

where *B* is the BLOSUM62 matrix and *B*_min_ is its global minimum. The penalty is 0 for the identity and 1 for the least favourable substitutions. Each position *i* carries a ground-truth importance *w*_*i*_ set by its annotation category (e.g. active, binding or metal-binding site); unannotated positions receive *w*_*i*_ = 0.05.

- **Additive:** each mutation lowers fitness by its weighted penalty,

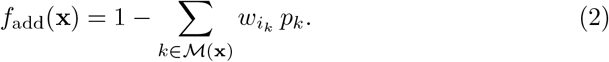

- **Epistatic:** the additive landscape is augmented with pairwise interactions between mutations,

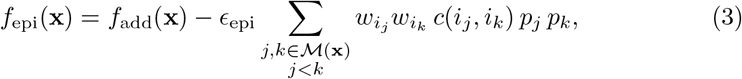

with *ϵ*_epi_ = 0.1. The contact proxy *c* is

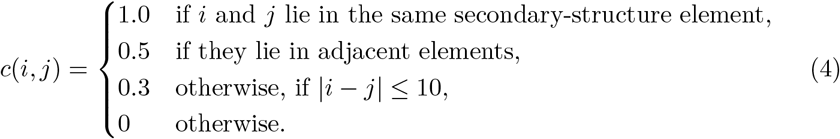

- **Nonlinear (stability cliff):** positions in each protein’s highest-weighted annotation category are designated critical (*C*). Mutations are split into a critical damage term and a non-critical loss term,

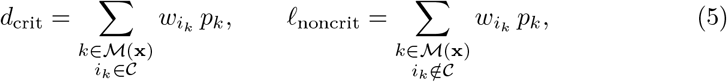

and fitness collapses once critical damage crosses a threshold:

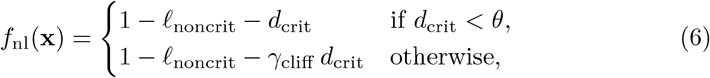

with threshold *θ* = 1.5 and cliff multiplier *γ*_cliff_ = 3. Non-critical positions therefore contribute additively regardless.

We sample 1, 500, 4, 000, and 10, 000 variants for GB1, T4 Lysozyme, and p53, respectively with 70% of the sampled variants being single mutants, 20% double mutants, and 10% triple mutants. An example walkthrough for how the synthetic datasets are built is presented in Appendix A Fig A.1.

#### 4.1.1 Label noise to perturb predictor accuracy

To obtain predictors spanning a range of accuracy on the same data, we corrupted the fitness labels by blending them with a random permutation of themselves. For a noise level *p* ∈ [0, 1], each label *y*_*j*_ was replaced by

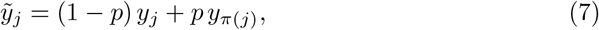

where *π* is a random permutation of the variants. The blended labels were then rescaled linearly to the mean and standard deviation of the original labels, so that both were unchanged at every *p*. Because *y*_*π*(*j*)_ is unrelated to *y*_*j*_, the correlation between noisy and true labels is approximately 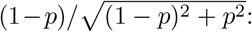 at *p* = 0 (exact labels), 0.71 at *p* = 0.5, and 0 at *p* = 1, where the labels are a full permutation and no sequence– fitness relationship remains to be learned. Unlike additive Gaussian noise, *p* does not depend on the scale of the labels, so the same levels are comparable across proteins and landscapes.

We used *p* ∈ {0, 0.05, 0.1, 0.2, 0.35, 0.5, 0.7, 0.85, 1}. Noise was applied once to all labels before the train/test split, with the permutation drawn from a fixed seed. In multi-seed experiments the noise seed was set equal to the training seed, so each seed sampled both a different initialization and a different noise realization. Because the test labels were corrupted too, test-set accuracy measures agreement with the noisy labels: even a predictor that recovered the true fitness exactly would correlate with the test labels only to the extent given above. Attributions were always scored against the noise-free ground truth of the synthetic landscape.

### 4.2 Protein language models and representation extraction

We represented each sequence by per-residue embeddings from a pretrained protein language model (pLM) whose weights were kept fixed throughout. We used three pLM families: ESM-C (300M & 600M parameters), Profluent E1 (150M, 300M, & 600M parameters) and Ankh-base (∼450M) Ankh-large ∼1.2B). For a sequence of length *L*, each model returns one *D*-dimensional vector per token. We removed the special tokens added by each tokenizer (start and end tokens for ESM-C and E1, the end token for Ankh), so that row *i* of the resulting *L × D* matrix corresponds to residue *i*. Embeddings were taken from the final layer, except in the representation-depth analysis, for which we extracted the hidden states of Ankh-base after layers 0 (the input embedding layer), 12, 24, 36 and 48 (the final layer). Embeddings were computed once per sequence and cached.

Before training, embeddings were standardized with a single scalar mean *µ* and standard deviation *σ* computed over all entries of the training-set embeddings, *x* ← (*x ™ µ*)*/σ*; test sequences were excluded from this calculation. The same *µ* and *σ* were stored with each trained predictor and reused when computing attributions, so that predictions and attributions were computed on identical inputs.

### 4.3 Predictor architectures

Every predictor maps the standardized embedding matrix *X* = (*x*_1_, …, *x*_*L*_) to a scalar fitness prediction 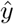. The four architectures differ in where the learned non-linear transformation sits relative to pooling, the step that combines residues into one sequence-level quantity. This placement determines how attribution can be distributed across residues, so we treated it as an experimental factor. All four share a four-layer perceptron whose widths were set per dataset; it uses GELU activations, dropout and a linear skip connection from the first to the fourth hidden layer.

### 4.4 Training and model selection

Each dataset was split once into a held-out test set (20% of variants) and a training set (80%) with a fixed split seed, so that all predictors for a dataset were evaluated on the same variants. Within the training set we ran five-fold cross-validation. In each fold, predictors were trained with AdamW (weight decay 10^−4^), a cosine learning-rate schedule decaying to 1% of the initial rate, Huber loss (*δ* = 1) and gradient-norm clipping at 1.0, for up to 50 epochs, with early stopping on the validation loss. The learning rate (5 × 10^−4^ to 10^−3^), batch size (64 to 256), dropout (0.3 to 0.5), early-stopping patience (15 to 30 epochs) and perceptron widths were set per protein and landscape. For a given dataset, all four architectures and all pLMs used the same settings. The final predictor was then trained on the full training set for the mean best epoch across folds (with a dataset-specific minimum of 10 to 20 epochs), with a linear learning-rate warm-up over the first tenth of training. Synthetic fitness values were used without rescaling (wild type = 1). Predictive accuracy was measured as Spearman’s *ρ* between predicted and observed fitness on the held-out test set. Each configuration was trained with 3-5 random seeds, depending on the experiment.

### 4.5 Integrated gradients attribution

We attributed each prediction to individual residues with integrated gradients (IG), using the wild type as the reference. Let **e**^0^ and **e**^1^ ∈ ℝ^*L×D*^ be the per-residue embeddings of the wild type and of a variant, and let *F* be the trained predictor. Along the straight-line path *γ*(*α*) = **e**^0^ + *α*(**e**^1^ − **e**^0^), the attribution to residue *i* is

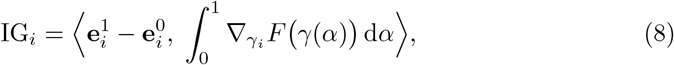

where the inner product runs over the *D* embedding dimensions, giving one signed value per residue, as visualized in Fig C.1. The attributions satisfy completeness, ∑*i*IG_*i*_ = *F* (**e**^1^) − *F* (**e**^0^), so they divide the predicted effect of the variant among its residues. Because pLM embeddings are contextual, 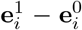, and hence IG_*i*_, can be non-zero at residues that were not mutated.

We approximated the integral with the trapezoidal rule on *S* = 500 uniform intervals and computed gradients by automatic differentiation through the embedding standardization, with dropout disabled. To limit discretization error, we first ran IG with 25 intervals and excluded variants whose completeness error, | Σ_*i*_ IG_*i*_ − (*F* (**e**^1^) − *F* (**e**^0^)), exceeded 0.1; attributions were then computed for all remaining training and test variants. More detailed definitions are provided in Appendix C.

### 4.6 Attribution metrics and TOPSIS aggregation

For each run, the per-residue attributions *A*_*i*_ of all analyzed sequences were summarized into two position scores and compared with the ground-truth importance *w*:

- **Magnitude score** *a*_i_ : the mean of |*A*_*i*_| plus the standard deviation of *A*_*i*_ .
- **Signed score** *s*_*i*_: the mean of *A*_*i*_.

Five metrics were computed:

- **Spearman correlation** between *a* and *w*.
- **Precision@***k*: the fraction of the *k* top-ranked positions in *a* that are annotated, where *k* is the number of annotated positions.
- **Top-category recall**: the fraction of the highest-weight annotated sites (at least five) that rank in the top *k* of *a*.
- **AUROC** for separating annotated from background positions using *a*.
- **Sign accuracy**, computed from *s*: the fraction of annotated positions with *s*_*i*_ *<* 0. Every substitution in the synthetic landscapes lowers fitness, so the correct sign is negative.

Scaling:

- Spearman correlation, precision@*k* and top-category recall were rescaled so that 0 is the mean value for random attributions and 1 is a perfect attribution, which makes them comparable across proteins with different annotation densities. The random mean was estimated from 400 random attribution vectors per protein.
- AUROC and sign accuracy were not rescaled in the TOPSIS computation; both equal 0.5 for random attributions.

We combined the five metrics with TOPSIS [38], using equal weights and computing scores separately for each combination of protein and landscape. TOPSIS builds an ideal point from the best observed value of each metric and an anti-ideal point from the worst, and scores every alternative by its relative distance to the two, from 0 at the anti-ideal to 1 at the ideal. Scores are therefore relative to the set of alternatives. Missing metric values were replaced by the worst value in the group. For the noise sweep (Fig 3) the alternatives were all ESM-C 300M runs, covering every architecture, comparator attribution, noise level and seed, plus the one-hot and shuffled-label controls. For the backbone comparison (Fig 4) they were pairs of predictor head and backbone, each at the noise level with its best mean held-out *ρ*; for the layer sweep, the Ankh-base layers; and for the ensemble analysis (Fig 5), bootstrap ensembles. We measured how closely attribution quality follows predictive accuracy as the Pearson correlation between TOPSIS score and held-out Spearman *ρ* over the multi-seed noise sweep, for each architecture and each combination of protein and landscape.

### 4.7 Cascading parameter randomization

To test whether attributions depend on the learned predictor parameters, we applied the cascading model-parameter randomization test of Adebayo et al. [32]. Starting from a trained predictor, we re-initialized its layers from the output toward the input, keeping earlier re-initializations in place, and recomputed the attributions after each step. For pool-first the order was: output layer; fourth hidden layer together with the skip projection; third, second and first hidden layers; layer normalization; attention-pooling layer. PreMLP has the same order followed by its per-residue network. In AttnPool the pooling-score layer is re-initialized immediately after the output layer, and MeanPool has no pooling parameters. Weights of linear and convolutional layers were redrawn from a Xavier-uniform distribution with zero biases, and layernormalization parameters were reset to unit scale and zero shift. The embedding standardization constants of the trained model were kept throughout.

At each step we computed linear-path IG (*S* = 100) for the same 50 held-out test variants, sampled once with a fixed seed, and averaged |IG_*i*_| over them to obtain one score per position. We report the Spearman correlation of this score with the score of the nonrandomized predictor and with *w*. The test was run on GB1 and T4 lysozyme (all three landscapes) at *p* = 0 for the four architectures, with three seeds each.

### 4.8 Ensembles

To reduce attribution variability, we formed ensembles of predictors trained with different seeds and averaged their position scores *a* and *s*. An ensemble consisted of predictors trained with the same data, train/test split, cross-validation folds, architecture and hyperparameters, differing only in the random seed. The seed set the weight initialization, the order of the mini-batches, the dropout masks and, when labels were noised, the permutation used to build the noisy labels. Attributions were computed separately for each member, and the ensemble attribution was the per-position average of the members’ magnitude scores *a* and signed scores *s*. Given *K* independently seeded models, each producing per-position scores 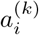 and 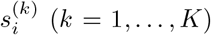, the ensemble attribution is the unweighted mean across seeds:

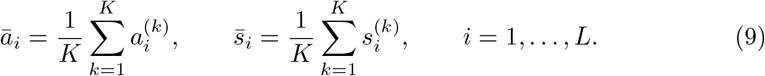

For the synthetic landscapes we trained up to three seeds for all reported results and ten seeds for the Fig 5, and five seeds for the real protein case studies. For the two real protein case studies, the five seeds of each configuration were averaged into consensus magnitude and signed scores per position, and agreement between seeds was summarized as the absolute mean signed score divided by its standard deviation across seeds.

### 4.9 Experimental datasets: *α*-amylase and *β*-lactamase

We evaluated IGLOO on two experimentally characterized enzyme families. For *α*-amylase, we used the FLIP2 *Bacillus α*-amylase site saturation mutagenesis library (3,706 variants of a 425-residue long wild type, quantitative activity measures the starch based stain removal by non-normalized secreted *α*-amylases), with all four of the benchmark’s prescribed train / test splits:

- `mutation/by_mutation`: (3,004 / 702)
- `number/one_to_many`: (3,218 / 488)
- `position/close_to_far`: (1,782 / 1,924)
- `position/far_to_close` (1,924 / 1,782)

Each split withholds a different method of extrapolation, so the four splits share a variant library but differ in the generalization demanded of the predictor. For *β*-lactamase, we used four ProteinGym TEM-1 deep mutational libraries built upon the following: Deng 2012 [39], Firnberg 2014 [40], Jacquier 2013 [41], and Stiffler 2015 [42]. All datasets carry a 286 residue wild type and range from 991–4,996 variants, each carrying its own train / test split. Assay values were *z*-scored within each dataset before training.

#### 4.9.1 Predictor and attribution extraction on experimental data

For every dataset, we trained the mean-pooling per-residue head on frozen Ankh-base embeddings taken from the final encoder layer. Heads were four-layer MLPs trained with the same training protocol described in Section 4.4 and Appendix B. Each dataset was trained with five independent seeds. Attributions were computed as described in Section 4.5, where embedding-level gradients along the straight-line path from the wild-type embedding to the variant embedding were used to give one signed value per residue position. We attributed every variant in each library rather than a selected subset, giving a position × variant attribution matrix per seed. For each position *i*, we averaged the signed attribution over all variants within a seed, then combined seeds into an ensemble mean 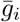, a cross-seed dispersion *sigma*_*i*_ (sample s.d. over the five seed means) and a reliability ratio:

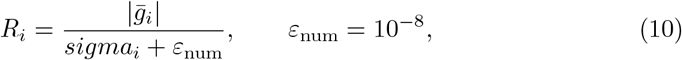

which is large when the five independently seeded models agree on both the magnitude and the sign of a position’s contribution. Cross-seed statistics were recomputed from the full-precision per-variant attribution tables rather than rounded per-position summaries, since rounding inflates *R*_*i*_ for small effects. Positions were then ranked by

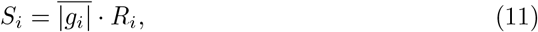

the ensemble mean absolute attribution weighted by its cross-seed reliability. An analogous substitution-level score was computed from the per-substitution attribution tables, 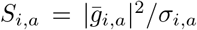, by averaging over all variants carrying substitution *a* at position *i*.

#### 4.9.2 Homolog search, multiple sequence alignment, and conservation scoring

Homologs of each wild-type sequence were retrieved with blastp on the NCBI server against the ClusteredNR database (*E ≤* 10^−5^, BLOSUM62 substitution matrix, word size 3). Hits were filtered to 35–95% sequence identity relative to reference to exclude near-duplicates and remote matches, then long sequences were trimmed to only retain the segment with the best BLAST scoring. The set was then clustered with MMseqs2 at 90% sequence identity to remove redundancy, aligned with MAFFT (L-INS-i), and each reference position was scored by 1 − *H*_norm_, where *H*_norm_ is the normalized Shannon entropy of its aligned column. A residue position with a score above 0.8 was defined as conserved.

#### 4.9.3 Rosetta Cartesian ΔΔ*G* prediction

Starting structures were stripped of all hetero atoms except for calcium co-factors in the case of *α*-amylase. Each structure was relaxed with unrestrained Cartesian FastRelax (4 repeats, 20 independent models, `fa_max_dis` = 9.0 Å), and the lowest-scoring model was used as the template for mutant model building. Mutational effects on stability were then computed over every residue position with one shared wild-type calculation and all 19 substitutions per position, 5 rounds each, `fa_max_dis` = 9.0 Å, `ddg_n_converged` = 2, `ddg_score_cutoff` = 1.0, `ddg_bbnbrs` = 1, `ddg_frag_nbrs` = 2 and `ddg_optimize_proline` disabled. For each substitution, ΔΔ*G* was taken as the lowest mutant score minus the lowest wild-type score. Reported ΔΔ*G* values are the mean ± sample standard deviation (*n* = 5) across five fully independent replicates of this protocol with REU converted to kcal mol^−1^ by dividing by 2.94. Positive values are defined as destabilizing.

#### 4.9.4 Functional categorization of *α*-amylase residue positions

Attributions were interpreted using a two-axis scheme that crosses the mutational effect of stability (ΔΔ*G*) with evolutionary constraint (conservation). The ΔΔ*G* axis was calibrated against measured activity rather than set by convention as follows: a two-component Gaussian mixture on the FLIP2 activity distribution labeled a variant impaired at function ≤ 0.0285, and across the 488 single mutants (99 impaired) the Youden-optimal ΔΔ*G* cutoff was 3.23 kcal mol^−1^ (ROC AUC 0.74, 95% CI 0.67–0.79), close to the peak-plus-2*σ* value of the ΔΔ*G* distribution itself (2.50 kcal mol^−1^) and well above the conventional 1 kcal mol^−1^ threshold, which flagged 59% of all substitutions. Since no ΔΔ*G* cutoff raised the precision for impaired variants much above 55%, we added conservation as a second, independent axis, with positions defined as conserved if they have a score above 0.8 (76 of 425 *α*-amylase positions). Crossing the two axes assigns every substitution to one of four categories:

- conserved and stability is sensitive to mutations (*structural + conserved*)
- conserved but stability is insensitive to mutations (*conserved*, i.e. constrained for catalysis, expression, folding, transport, etc.)
- not conserved but stability is sensitive to mutations (*structural*)
- neither (*background*)

This partitions the 425 × 19 = 8,075 substitutions into 677 structural (8%), 767 catalytic/signaling (9%), 1,639 detergent-environment (20%) and 4,992 background (62%) for *α*-amylase. As a check on the scheme, the putative catalytic triad (D176, E208, D269) is ∼99% conserved with ΔΔ*G ≈* 0 and falls almost entirely in the catalytic/signaling category.

Category enrichment among top-ranked attributions was assessed against all substitutions observed in the library, with significance from a position-level permutation test (20,000 draws of *K* random positions) and a bootstrap over positions (2,000 resamples), so that substitutions sharing a position are not treated as independent observations. Finally, the positions identified by each split were compared with published annotations of residue function.

#### 4.9.5 Selection of TEM-1 mutant candidates for follow-up analysis

Within each split, we ranked positions by *S*_*i*_ (Eq. 11) and retained the top five. At each retained position, we inspected the per-substitution attributions and selected those whose magnitude was clearly separated from the remaining substitutions at that position, yielding a short list of substitutions that the predictor singles out rather than a site directed mutagenesis scan. These candidates were carried forward to the same Rosetta cartesian ΔΔ*G* prediction as described prior and Boltz-2 ligand binding affinity prediction.

#### 4.9.6 Boltz-2 complex prediction

Each selected mutant was modeled with Boltz-2 using the model’s default multiple-sequence-alignment settings, with inference run on the mutant sequence together with the ligand associated with that dataset’s assay. Reported affinity values are the mean ± sample standard deviation (*n* = 20) across twenty independent Boltz-2 runs.

### 4.10 Compute

All experiments were run on HPC clusters. Downstream predictor heads are small enough that most jobs requested a single NVIDIA H100 MIG partition (1g.10gb, i.e. 1/7 of a physical GPU) rather than a full GPU; training a single predictor head took ∼10 minutes. The full synthetic benchmark sweep (architectures × backbones × landscapes × noise levels × seeds) consumed approximately 1,170 full-GPU-equivalent hours.

## Acknowledgements

We thank Ahmed Sallam for helpful discussions on the interpretability of learned representations in other biological foundational models and the application of IGLOO to other domains.

## Data and Code Availability

The synthetic protein datasets as well as the code is all available in https://github.com/chemcognition-lab/IGLOO.

## Author Contribution

Conceptualization: R.B., B.S.L. Methodology: R.B. Investigation: R.B., S.W., J.M.B.C., W.C. Formal analysis: R.B., W.C. Data curation: R.B., S.W. Software: R.B., S.W., J.M.B.C. Validation: P.D. Writing – original draft: R.B., S.W., J.M.B.C., W.C. Writing – review & editing: all authors. Supervision: K.M., B.S.L. Funding acquisition: K.M., B.S.L.

## Generative AI Tools Disclosure

Generative AI tools (LLMs) were used during the preparation of this manuscript to assist with text editing, code generation, and language polishing. ChatGPT and Claude were used to refine the clarity, grammar, and flow of the initial draft. Claude was used to assist in writing and debugging the data analysis and model training scripts. The authors reviewed and edited all AI-generated outputs and take full responsibility for the final text, code, and accuracy of the scientific content.

## Appendix A Synthetic Protein Datasets

Here, we present an example walkthrough of the GB1 protein and how the synthetic fitness landscapes are built for a double mutant. It is presented in Fig A.1.

**Fig. A.1.**
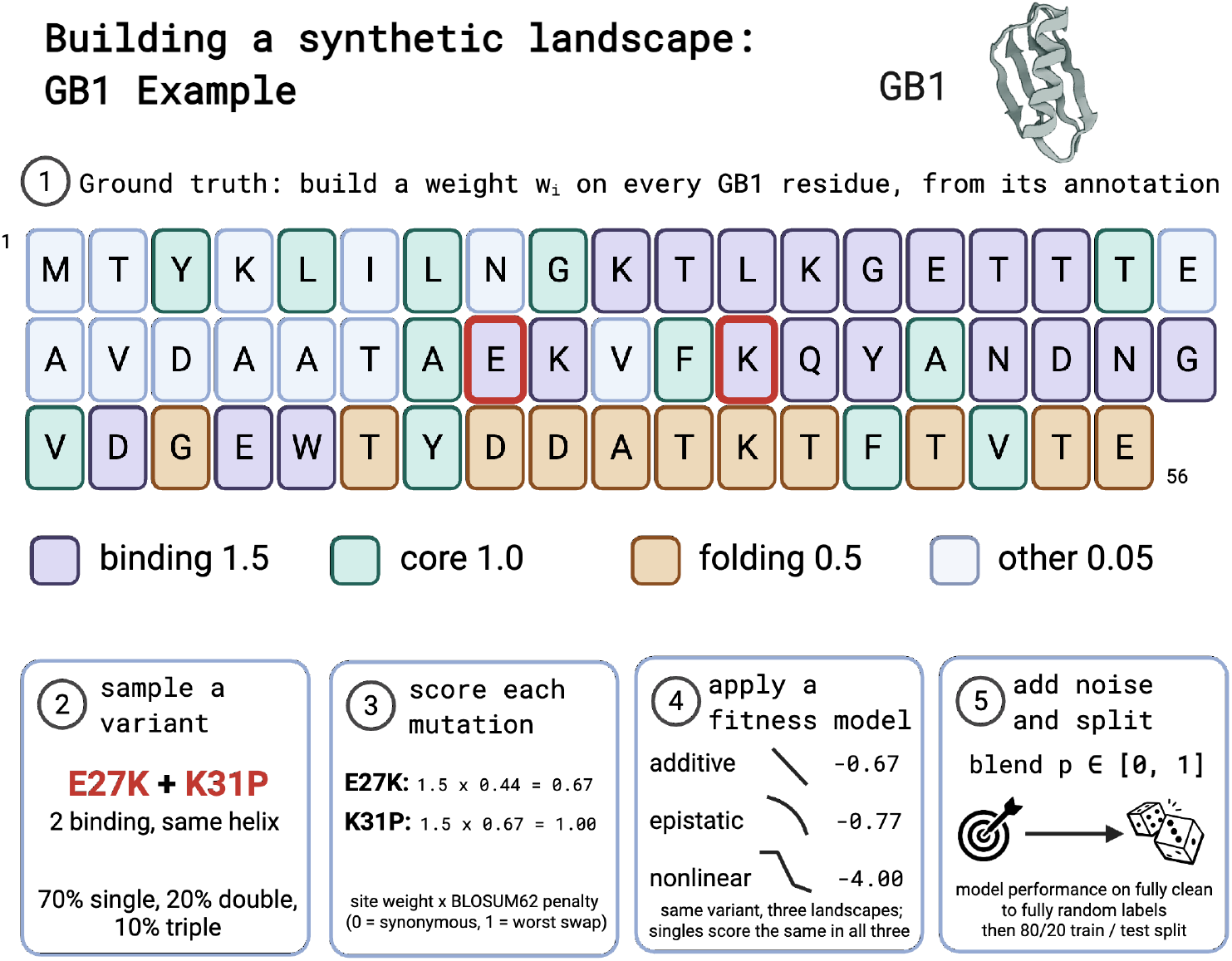
GB1 synthetic data example walkthrough. **1** We build a ground-truth per-residue attribution by assigning weights from experimentally reported functional sites. **2** Next, we sample variants by randomly mutating the wild-type sequence. Here, we use the E27K and K31P doublemutant as an example. Both of these mutations lie on binding sites and on the same helix. **3** We then score each mutation by multiplying the assigned weight of the mutated position with the BLOSUM62 matrix amino acid substitution penalty. **4** We apply the 3 different fitness models independently so each mutation is assigned three different fitness values. The additive landscape linearly combines the mutation scores; the epistatic landscape adds a higher contact penalty for mutations that are structurally close; the nonlinear landscape maintains an additive effect for mutants until a damage threshold is reached, and the fitness value drops in a cliff-like manner. **5** Finally, we add noise to the fitness labels to test model performance ranging from fully clean labels to fully randomized labels.

### A.1 Rationale for Synthetic Landscapes

We preface this section by addressing the motivation for not using biological data for experimental work. Real biological ground truth for positional attribution is intractable at scale due to the different attributions to different protein properties by the same amino acid at each residue position. As such, multiple repeats of deep mutational scanning on target proteins to collect data to assign attribution scores for different properties become a giant undertaking. As such, assignment of importance scores by researchers will vary based on research objective and heuristics. By employing a synthetic landscape design, we can leverage computable ground truths to systematically isolate accuracy, stability, and fidelity.

**Table A.1.** Summary of the three benchmark proteins used in IGLOO evaluation. Annotation density is the fraction of positions with importance weight *w*_*i*_ *> w*_bg_. Positions are reported using 1-indexed residue numbering.

| Protein | UniProt | Length | Ann. sites | Ann. density |
| --- | --- | --- | --- | --- |
| GB1<br>IgG-binding B1<br>domain | P06654 | 56 | 43 | 77% |
| Binding-site ($w = 1.5$): Fc-interface (E27, K28, K31, Q32), Fab-interface (K10–T17, Y33, N35–G38) | | | | |
| T4 Lysozyme<br>Bacteriophage<br>lytic enzyme | P00720 | 164 | 7 | 4% |
| Active-site ($w = 5.0$): Glu11, Asp20 (catalytic dyad) | | | | |
| p53<br>Human tumour<br>suppressor | P04637 | 393 | 41 | 10% |
| Metal-binding ($w = 5.0$): C176, H179, C238, C242 ($Zn^{2+}$ coordination tetrad) | | | | |

Before computing any metric, sequences with |*f*| *< δ*_filter_ (default *δ*_filter_ = 0.01) are removed by a *model collapse filter*: variants with near-zero fitness effects carry no positional signal and would dilute all position-level measures.

## Appendix B Model Architecture & Training

In the baseline fitness models, pLM sequence embeddings are extracted and aggregated using attention-weighted pooling, then passed through a four-layer MLP with ReLU activations, Huber loss, and dropout regularization. Training uses Adam with a cosine-annealed learning rate, 5-fold cross-validation with an 80/20 train–test split, and early stopping with a minimum-epoch floor. Integrated gradients are computed with 500 interpolation steps. Hyperparameters that are shared across all models are summarized in Table B.1; per-model choices are given in Table B.4.

We tested alternative architectures that shuffle the pooling & processing heads in the model. The model architectures are described below.

### B.1 Tested Models & Architectures

#### B.1.1 OneHot

This baseline removes the pLM. Each residue is a one-hot vector of its amino-acid identity (the 20 standard amino acids plus one channel for any other symbol, *D* = 21), so *x*_*i*_ carries no information about neighboring residues or position. Standardization, training and integrated gradients are otherwise unchanged, and the head is pool-first (216,705 trainable parameters for GB1 with widths 512, 256, 128 and 64, against 700,290 with ESM-C embeddings). The baseline separates the contribution of the pLM from that of amino-acid identity alone. For a variant that differs from the wild type only by substitutions, 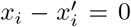 at every unmutated residue, so IG assigns those residues exactly zero attribution; with pLM embeddings, a substitution also changes the embeddings of other residues, and attribution can land on residues that were not mutated. Because the attention score *t*_*i*_ = *w*^⊤^*x*_*i*_ depends only on residue identity, this head cannot distinguish positions and acts on an attention-weighted amino-acid composition.

#### B.1.2 Pool-First

This is the FLIP-style attention-pooling baseline. A scalar score *t*_*i*_ = *w*^⊤^*x*_*i*_ + *b* is computed independently at each position (a width-1 convolution), normalized across positions as 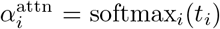, and used to pool the embeddings, 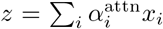. The pooled vector passes through layer normalization and the four-layer perceptron, and a final linear layer outputs 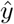. For GB1 with ESM-C embeddings (widths 512, 256, 128 and 64) the head has 700,290 trainable parameters. Because every learned nonlinearity acts after pooling, the gradient at residue *i* is, apart from the dependence of 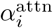 on *x*_*i*_, the gradient at the pooled vector scaled by the attention weight, 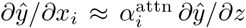. The attention weights, therefore, largely set the attribution pattern.

#### B.1.3 PreMLP

This architecture is identical to pool-first, except that each embedding first passes through a residual network shared across positions, *x*_*i*_ *← x*_*i*_ + *g*(*x*_*i*_), where *g* is a two-layer perceptron of width *D* with a GELU activation and dropout. Pooling and the downstream head are unchanged, so the comparison with pool-first isolates the effect of a learnable per-residue transformation before pooling.

#### B.1.4 AttnPool

The whole perceptron is moved before pooling and applied to each residue with shared weights. Each embedding is layer-normalized and passed through the four-layer perceptron, giving a hidden representation *h*_*i*_; a linear readout gives a scalar contribution *f*_*i*_ = *v*^⊤^*h*_*i*_ + *c*. Attention weights are computed from the same hidden representation, 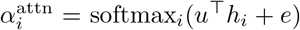, and the prediction is the weighted sum 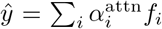. Pooling is the last operation, so each residue’s contribution is computed by a trained function of that residue before residues are combined.

#### B.1.5 MeanPool

The per-residue network is the same as in AttnPool, but the contributions *f*_*i*_ = *f* (*x*_*i*_) are combined by an unweighted mean, *ŷ*= *L*^−1^ Σ_*i*_ *f* (*x*_*i*_), with no learned pooling weights. Because the prediction is a sum of terms that each depend on a single residue, the IG attribution of residue *i*, summed over embedding dimensions, equals that residue’s own term, 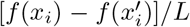, where 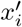 is the wild-type embedding at position *i*.

### B.2 Attribution Method Baselines

In addition to the architecture variants above, we compared IG against four alternative ways of extracting a per-residue attribution from a trained predictor, with the architecture (and hence *F*) held fixed. All baselines operate on the same wild-type and variant embeddings **e**^0^, **e**^1^ used by IG (Section 4.5).

#### B.1.2 ISM (in-silico mutagenesis)

A gradient-free, model-dependent reference. For each position *i*, every one of the 19 non-wild-type amino acids is substituted in turn and the trained predictor is reevaluated on the resulting sequence:

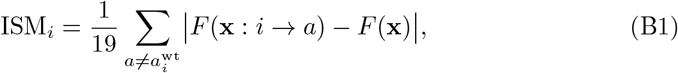

where **x** : *i* → *a* denotes the sequence with position *i* substituted to amino acid *a*. Unlike IG, ISM requires no gradients or embedding-space assumptions, at the cost of 19*L* forward passes per sequence.

#### B.2.2 Gradient Magnitude

Drops IG’s embedding-displacement weighting, keeping only the magnitude of the path-averaged gradient:

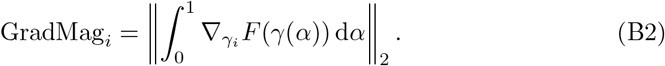

This isolates how sensitive the model is to position *i*’s embedding, independent of how much the variant actually displaced it.

#### B.2.3 Input Displacement

A model-independent control that discards the predictor’s gradient entirely, retaining only the geometry of the embedding change:

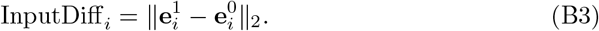

Because it does not depend on *F*, this baseline is identical across seeds, noise levels, and model quality for a given variant, and isolates how much of IG’s signal is explained by pLM embedding geometry rather than by anything the predictor learned.

### B.24 Gradient × Input

A single-point variant of IG that evaluates the gradient only at the endpoint of the path (*α* = 1), with no integration and no reference baseline:

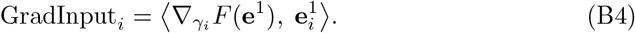

On the MeanPool and AttnPool architectures paired with Ankh-base embeddings, this baseline is degenerate: the single-point gradient saturates against the predictor’s output range, giving an undefined Spearman correlation and sign accuracy fixed at ™1 across every protein, landscape, and seed. We therefore exclude it from those two comparisons while retaining it for pool-first, where it behaves normally.

**Table B.1.**
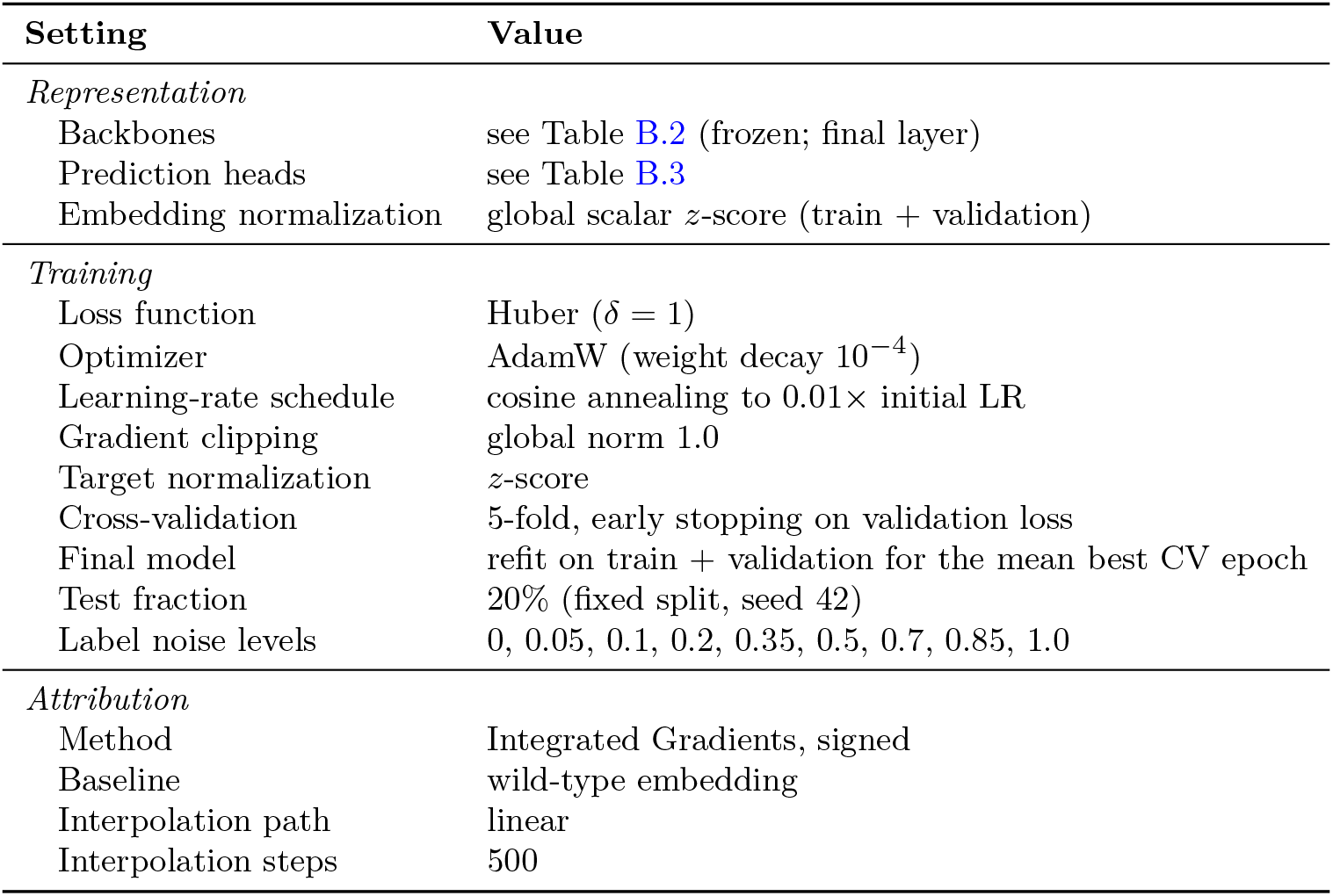
Settings shared across all synthetic-data experiments.

#### Model Test-Set Performance

Table B.5 reports held-out test-set performance (20% split, *n* = 289–1,902 sequences) of the final trained model at zero label noise for each protein–fitness-model combination.

**Table B.2.** Protein representations. All pLMs are used frozen, taking per-residue embeddings from the final layer with special tokens removed.

| Family | Model | Embedding dim. $D$ |
| --- | --- | --- |
| ESM-C | ESM-C 300M | 960 |
|  | ESM-C 600M | 1152 |
| Ankh | Ankh-base | 768 |
|  | Ankh-large | 1536 |
| Profluent E1 | E1 150M | 768 |
|  | E1 300M | 1024 |
|  | E1 600M | 1280 |
| None | One-hot | 21 |

**Table B.3.** Prediction heads. “MLP” is the four-layer GELU network with widths *d*_1_–*d*_4_ (Table B.4), dropout, and a residual connection from layer 1 to layer 4; it is shared across positions in the per-residue heads.

| Head | Structure |
| --- | --- |
| Pool-First | attention pooling over residues → LayerNorm → MLP → fitness |
| PreMLP | residual per-residue MLP ($D \rightarrow D \rightarrow D$) → Pool-First head |
| MeanPool | per-residue LayerNorm → MLP → per-residue score; fitness = mean over residues |
| AttnPool | as MeanPool, but fitness = attention-weighted sum of per-residue scores (weights from the MLP’s last hidden layer) |

**Table B.4.**
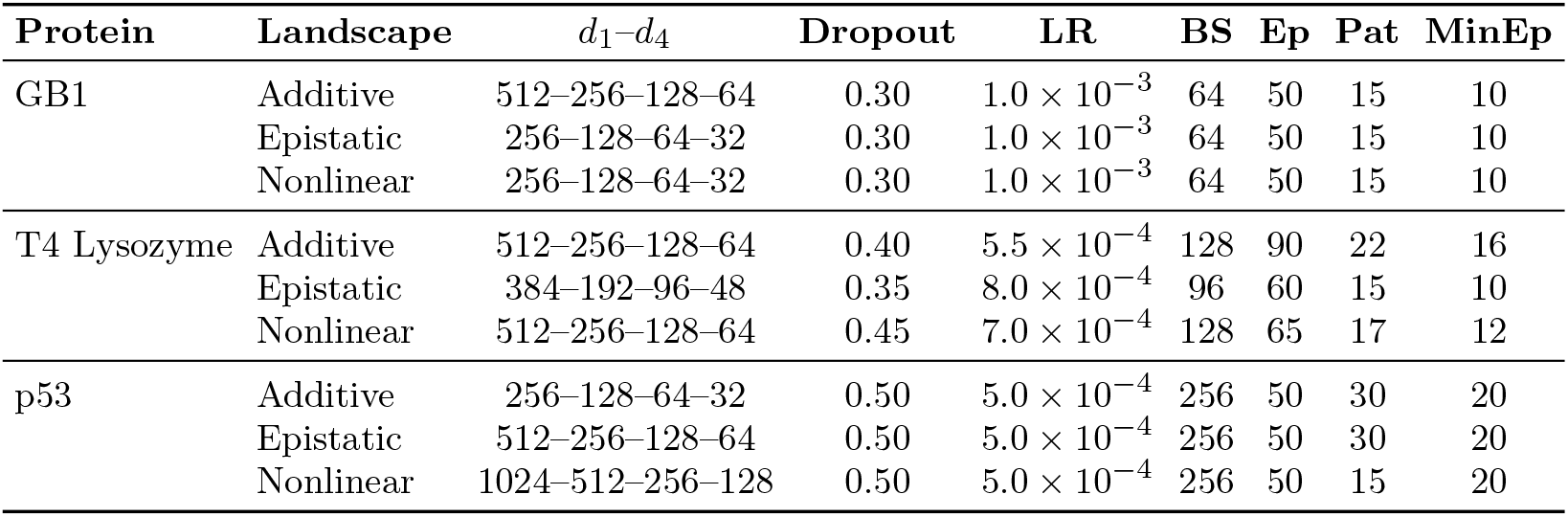
Per-landscape training hyperparameters, used for every backbone and prediction head. *d*_1_–*d*_4_: MLP hidden widths; LR: initial learning rate; BS: batch size; Ep: maximum epochs per CV fold; Pat: early-stopping patience; MinEp: minimum number of epochs for the final refit.

**Table B.5.**
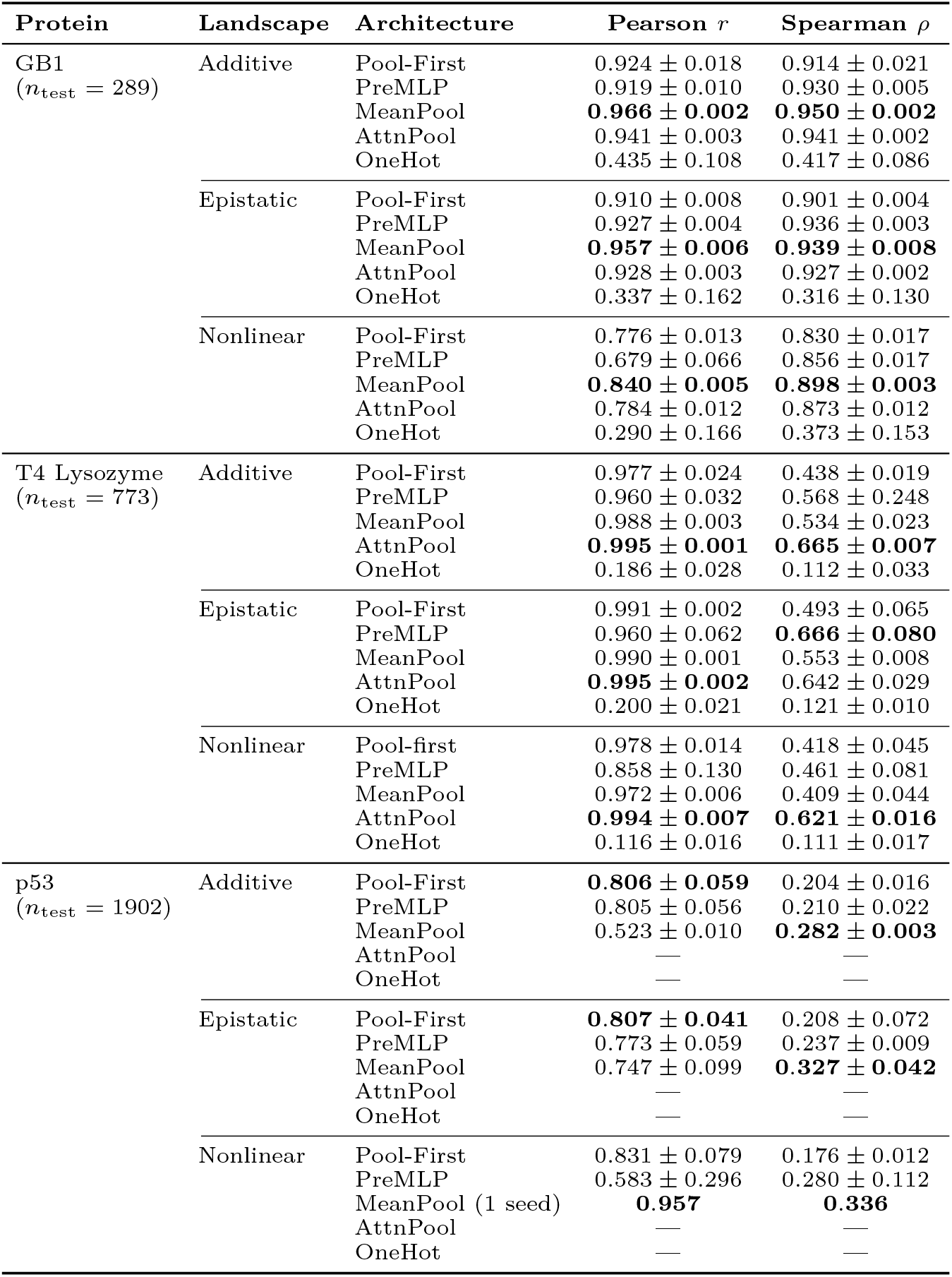
Predictor performance on the held-out test set (label noise = 0.0). Values are mean ± s.d. across seeds 1, 2 and 3; cells with fewer seeds note the count. The best mean per landscape is shown in bold. All architectures use the ESM-C 300M backbone except OneHot, which applies the Pool-First head to one-hot encodings. — indicates no runs.

| Protein | Landscape | Architecture | Pearson $r$ | Spearman $\rho$ |
| --- | --- | --- | --- | --- |
| GB1<br>($n_{\text{test}} = 289$) | Additive | Pool-First | $0.924 \pm 0.018$ | $0.914 \pm 0.021$ |
| | | PreMLP | $0.919 \pm 0.010$ | $0.930 \pm 0.005$ |
|  |  | MeanPool | **$0.966 \pm 0.002$** | **$0.950 \pm 0.002$** |
| | | AttnPool | $0.941 \pm 0.003$ | $0.941 \pm 0.002$ |
| | | OneHot | $0.435 \pm 0.108$ | $0.417 \pm 0.086$ |
| | Epistatic | Pool-First | $0.910 \pm 0.008$ | $0.901 \pm 0.004$ |
| | | PreMLP | $0.927 \pm 0.004$ | $0.936 \pm 0.003$ |
|  |  | MeanPool | **$0.957 \pm 0.006$** | **$0.939 \pm 0.008$** |
| | | AttnPool | $0.928 \pm 0.003$ | $0.927 \pm 0.002$ |
| | | OneHot | $0.337 \pm 0.162$ | $0.316 \pm 0.130$ |
| | Nonlinear | Pool-First | $0.776 \pm 0.013$ | $0.830 \pm 0.017$ |
| | | PreMLP | $0.679 \pm 0.066$ | $0.856 \pm 0.017$ |
|  |  | MeanPool | **$0.840 \pm 0.005$** | **$0.898 \pm 0.003$** |
| | | AttnPool | $0.784 \pm 0.012$ | $0.873 \pm 0.012$ |
| | | OneHot | $0.290 \pm 0.166$ | $0.373 \pm 0.153$ |
| T4 Lysozyme<br>($n_{\text{test}} = 773$) | Additive | Pool-First | $0.977 \pm 0.024$ | $0.438 \pm 0.019$ |
| | | PreMLP | $0.960 \pm 0.032$ | $0.568 \pm 0.248$ |
| | | MeanPool | $0.988 \pm 0.003$ | $0.534 \pm 0.023$ |
|  |  | AttnPool | **$0.995 \pm 0.001$** | **$0.665 \pm 0.007$** |
| | | OneHot | $0.186 \pm 0.028$ | $0.112 \pm 0.033$ |
| | Epistatic | Pool-First | $0.991 \pm 0.002$ | $0.493 \pm 0.065$ |
| | | PreMLP | $0.960 \pm 0.062$ | **$0.666 \pm 0.080$** |
| | | MeanPool | $0.990 \pm 0.001$ | $0.553 \pm 0.008$ |
| | | AttnPool | **$0.995 \pm 0.002$** | $0.642 \pm 0.029$ |
| | | OneHot | $0.200 \pm 0.021$ | $0.121 \pm 0.010$ |
| | Nonlinear | Pool-first | $0.978 \pm 0.014$ | $0.418 \pm 0.045$ |
| | | PreMLP | $0.858 \pm 0.130$ | $0.461 \pm 0.081$ |
| | | MeanPool | $0.972 \pm 0.006$ | $0.409 \pm 0.044$ |
|  |  | AttnPool | **$0.994 \pm 0.007$** | **$0.621 \pm 0.016$** |
| | | OneHot | $0.116 \pm 0.016$ | $0.111 \pm 0.017$ |
| p53<br>($n_{\text{test}} = 1902$) | Additive | Pool-First | **$0.806 \pm 0.059$** | $0.204 \pm 0.016$ |
| | | PreMLP | $0.805 \pm 0.056$ | $0.210 \pm 0.022$ |
| | | MeanPool | $0.523 \pm 0.010$ | **$0.282 \pm 0.003$** |
|  |  | AttnPool | — | — |
|  |  | OneHot | — | — |
| | Epistatic | Pool-First | **$0.807 \pm 0.041$** | $0.208 \pm 0.072$ |
| | | PreMLP | $0.773 \pm 0.059$ | $0.237 \pm 0.009$ |
| | | MeanPool | $0.747 \pm 0.099$ | **$0.327 \pm 0.042$** |
|  |  | AttnPool | — | — |
|  |  | OneHot | — | — |
| | Nonlinear | Pool-First | $0.831 \pm 0.079$ | $0.176 \pm 0.012$ |
| | | PreMLP | $0.583 \pm 0.296$ | $0.280 \pm 0.112$ |
|  |  | MeanPool (1 seed) | **$0.957$** | **$0.336$** |
|  |  | AttnPool | — | — |
|  |  | OneHot | — | — |

## Appendix C Integrated Gradients

We used integrated gradients (IG) (Fig C.1) because it naturally fits the structure of variant-effect prediction. IG has a built-in baseline in the wild-type sequence: every variant is defined relative to it, so attributions answer a biologically meaningful question, which is what residues account for the predicted change in fitness from wild type. IG satisfies completeness, so per-residue attributions sum to *F* (variant) − *F* (WT) and are expressed in units of fitness. It also satisfies implementation invariance: attributions depend only on the learned input–output function, not on how that function is parameterized. Differences in attribution between prediction heads, therefore, reflect genuine differences in the functions they learn, including along the interpolation path, rather than artifacts of the attribution method. Local gradient methods such as saliency and gradient × input rely on the gradient at a single point. They can therefore miss saturated or nonlinear effects of the kind built into our epistatic and nonlinear landscapes. Perturbation approaches such as occlusion or embedding ablation require replacing pLM embeddings with reference values that may lie outside the distribution the model was trained on. Shapley-value methods are intractable at the scale of *L × D* embedding inputs. IG needs only gradient access to a frozen-embedding predictor and a fixed number of model evaluations, making it practical across thousands of variants and many trained models. We report gradient × input, input-difference, and in-silico mutagenesis as baselines for comparison.

### C.1 Interpolation Methods: Full Definitions

All interpolation methods operate in the pLM embedding space. For a sequence variant *v* and the wild-type (WT) baseline *v*_WT_, let **e**^0^ ∈ ℝ^*L×D*^ and **e**^1^ ∈ ℝ^*L×D*^ denote the per-residue embeddings of *v*_WT_ and *v*, respectively, extracted from the frozen pLM, where *L* is the number of residues (special tokens such as BOS/EOS are removed) and *D* is the embedding dimension. We write **e**_*i*_ ∈ ℝ^*D*^ for the embedding of residue *i*. The integrated gradients attribution for position *i* is the scalar

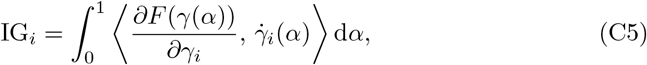

where *F* is the trained predictor, *γ*(*α*) ∈ ℝ^*L×D*^ is the interpolation path with *γ*(0) = **e**^0^ and 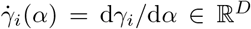 is the path tangent at position *i*, and ⟨·,·⟩ denotes the inner product over the embedding dimension. For the linear and slerp paths, the integral is approximated with the trapezoidal rule on *S* uniform intervals (*α*_*k*_ = *k/S, k* = 0, …, *S*).

The predictor is applied as 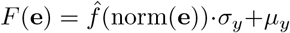. Here norm(**e**) = (**e** − *µ*_*e*_)*/σ*_*e*_ is a global normalization in which *µ*_*e*_ and *σ*_*e*_ are a single scalar mean and standard deviation computed over all entries of the training and validation embeddings; 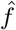 is the regression head; and (*µ*_*y*_, *σ*_*y*_) are the constants used to z-score the training targets. Gradients are taken with respect to the unnormalized embeddings through the full composition norm 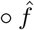, so the attributions reflect the complete computational graph.

**Fig. C.1.**
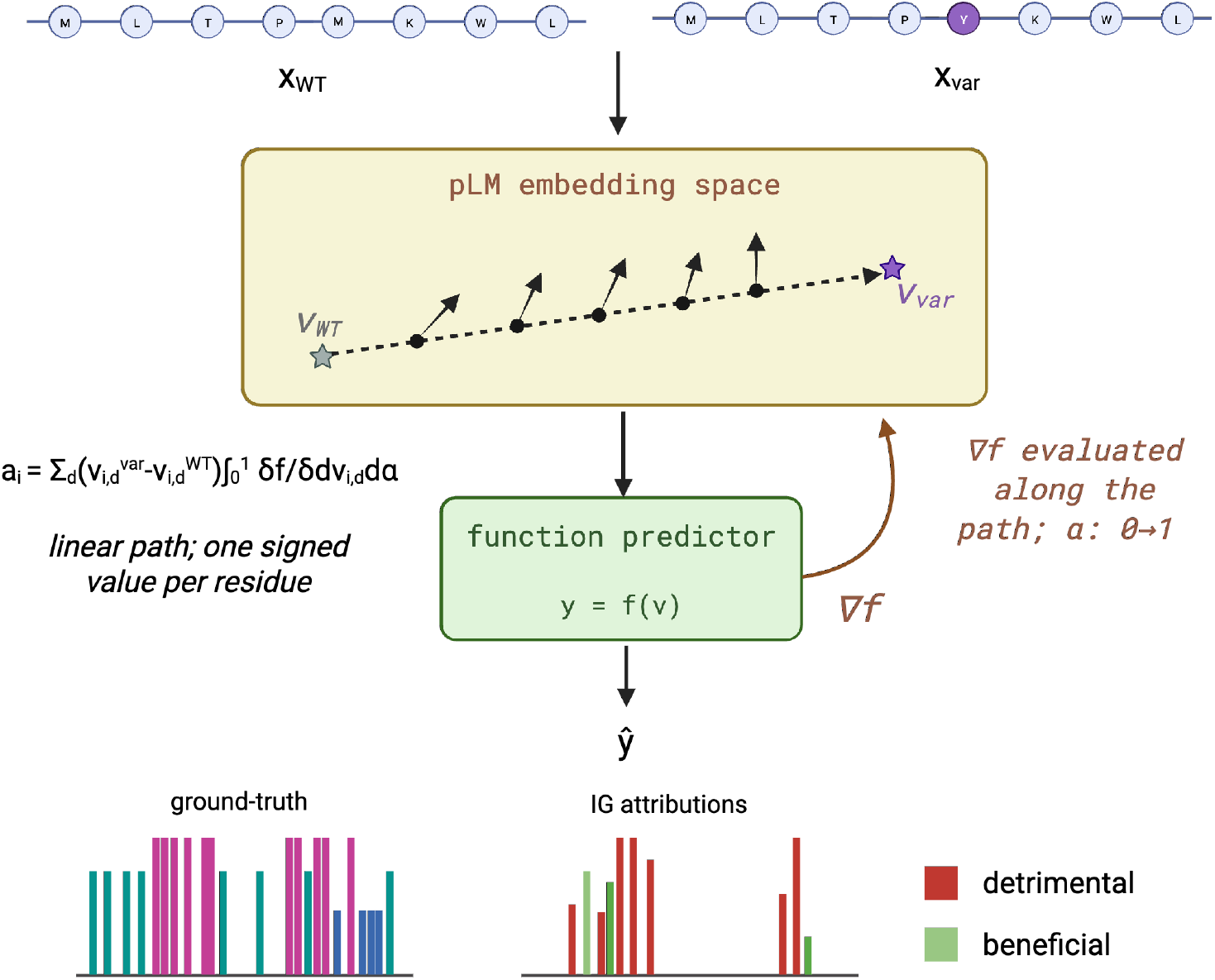
Integrated gradients in the embedding space for residue-level attributions.

#### Linear Interpolation

The linear path is the standard IG path [6]:

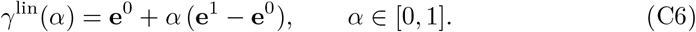

Its tangent is constant, 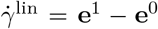, so the trapezoidal approximation of Eq. C5 reduces to the classic IG estimator

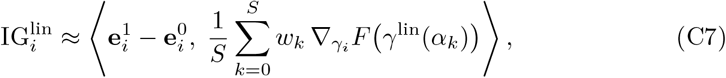

where *w*_0_ = *w*_*S*_ = 1*/*2 and *w*_*k*_ = 1 for 0 *< k < S* are the trapezoidal weights. The signed attributions satisfy completeness up to discretization error: 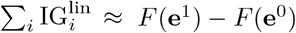

The *S* + 1 gradient evaluations along the linear path are independent, so they are computed in parallel mini-batches. The linear path also needs no extra computation to build the path or its tangent, which makes it the simplest method to compute.

### C.2 Integrated Gradients: Full Definitions

We used IG to assign each residue a share of the difference between the predicted fitness of a variant and that of the wild type. Let *F* denote a trained predictor, which returns predicted fitness, and let *X* and *X*^*′*^ in ℝ ^*L×D*^ be the embedding matrices of a variant and of the wild-type sequence, computed with the same pLM and layer. All variants are substitutions, so *X* and *X*^*′*^ have the same length and their rows correspond residue by residue. The wild type serves as the reference input (the IG baseline), so the quantity being explained is the predicted effect of the variant’s mutations, Δ*F* = *F* (*X*) − *F* (*X*^*′*^).

IG moves the input from the reference to the variant along a path and accumulates how fast the prediction changes along the way. We used the straight-line path

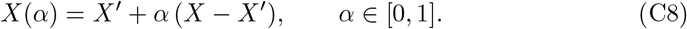

Along this path, the rate of change of the prediction is the directional derivative of *F* in the direction *X* − *X*^*′*^,

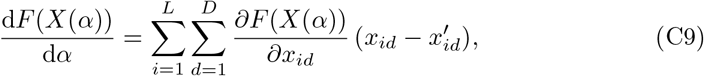

and integrating this rate over *α* recovers the total change in the prediction, 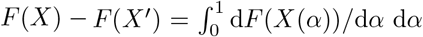, by the fundamental theorem of calculus for line integrals. Splitting the integral by input feature gives the IG attribution of embedding dimension *d* at residue *i*,

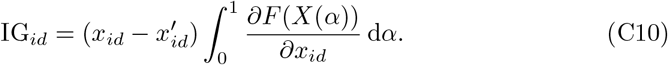

By construction, the attributions sum to the predicted effect, ∑_i,d_ IG_*id*_ = *F* (*X*) − *F* (*X*^*′*^) (completeness), so every attribution is expressed in units of fitness. IG can be defined along other paths between *X*^*′*^ and *X*; we used the straight line, the path of the original formulation and the only path method that treats symmetric input features identically.

### C.2.1 Per-residue attribution

We summed attributions over embedding dimensions to obtain one signed score per residue,

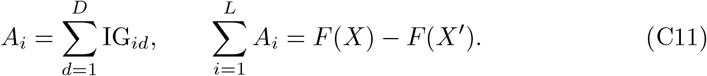

A positive *A*_*i*_ means that the change at residue *i* relative to the wild type raised the predicted fitness; a negative *A*_*i*_ means it lowered it. Because pLM embeddings are contextual, a substitution at one residue can change the embeddings of other residues, so 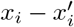, and therefore *A*_*i*_, can be non-zero at residues that were not mutated. The aggregation method for per-residue scores across variants and their comparison with the ground truth is described under Attribution metrics (Section 4.6).

#### Numerical integration

We approximated the integral with a Riemann sum over *m* points along the path, *α*_*k*_ = *k/m* for *k* = 1, …, *m*, averaging the gradients at those points:

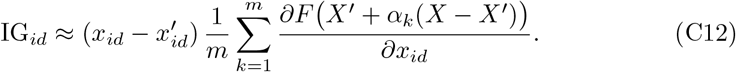

Gradients were computed by automatic differentiation of *F* with respect to the input embeddings, with the predictor in evaluation mode so that dropout was disabled. The path was interpolated in the original embedding space, and the standardization described above was applied inside the forward pass. Because the standardization is affine, the resulting attributions are identical to those computed directly on standardized embeddings.

## Appendix D Synthetic Data Results

**Fig. D.1.**
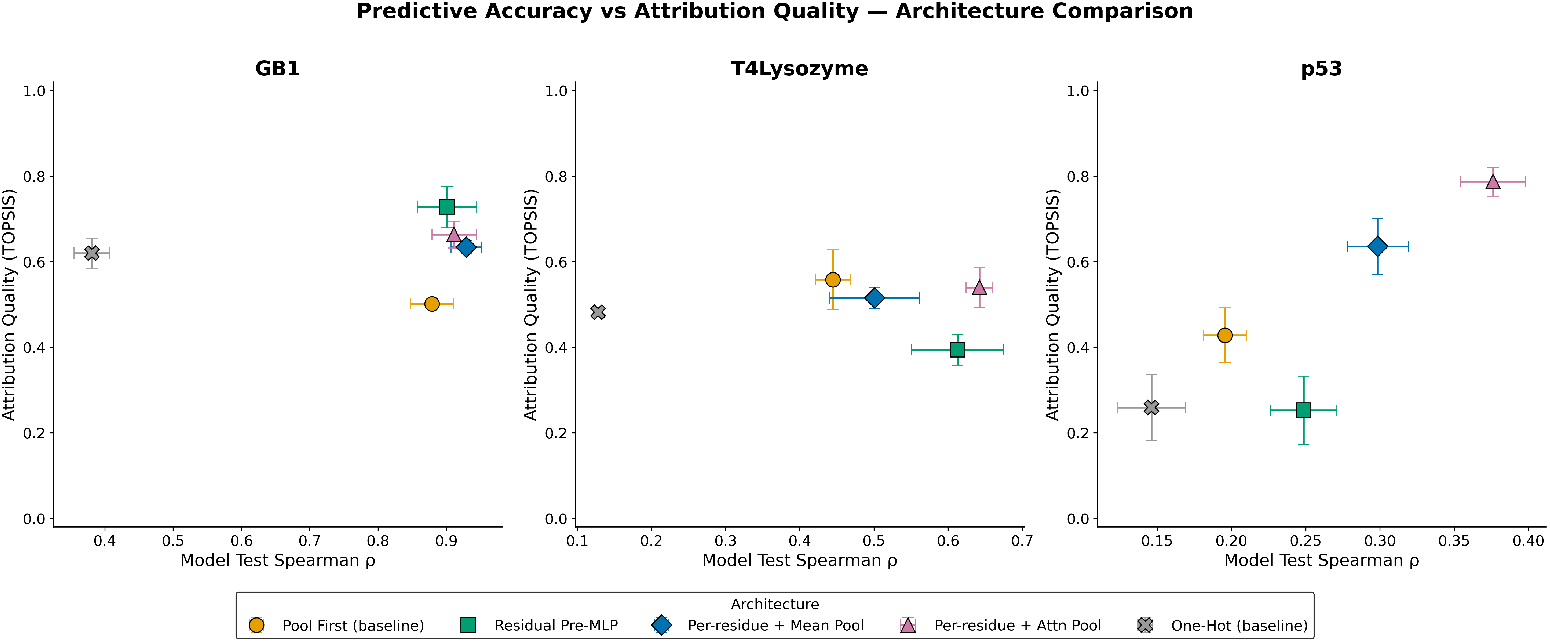
Predictive accuracy vs. attribution quality across different architectures. TOPSIS score (y-axis) was aggregated across the different landscapes for each protein and compared against the model test Spearman performance (x-axis). ESM-C 300M was used here across all the architectures (excluding the one-hot baseline) and the label noise with the best model test performance was chosen for comparison in this figure.

**Fig. D.2.**
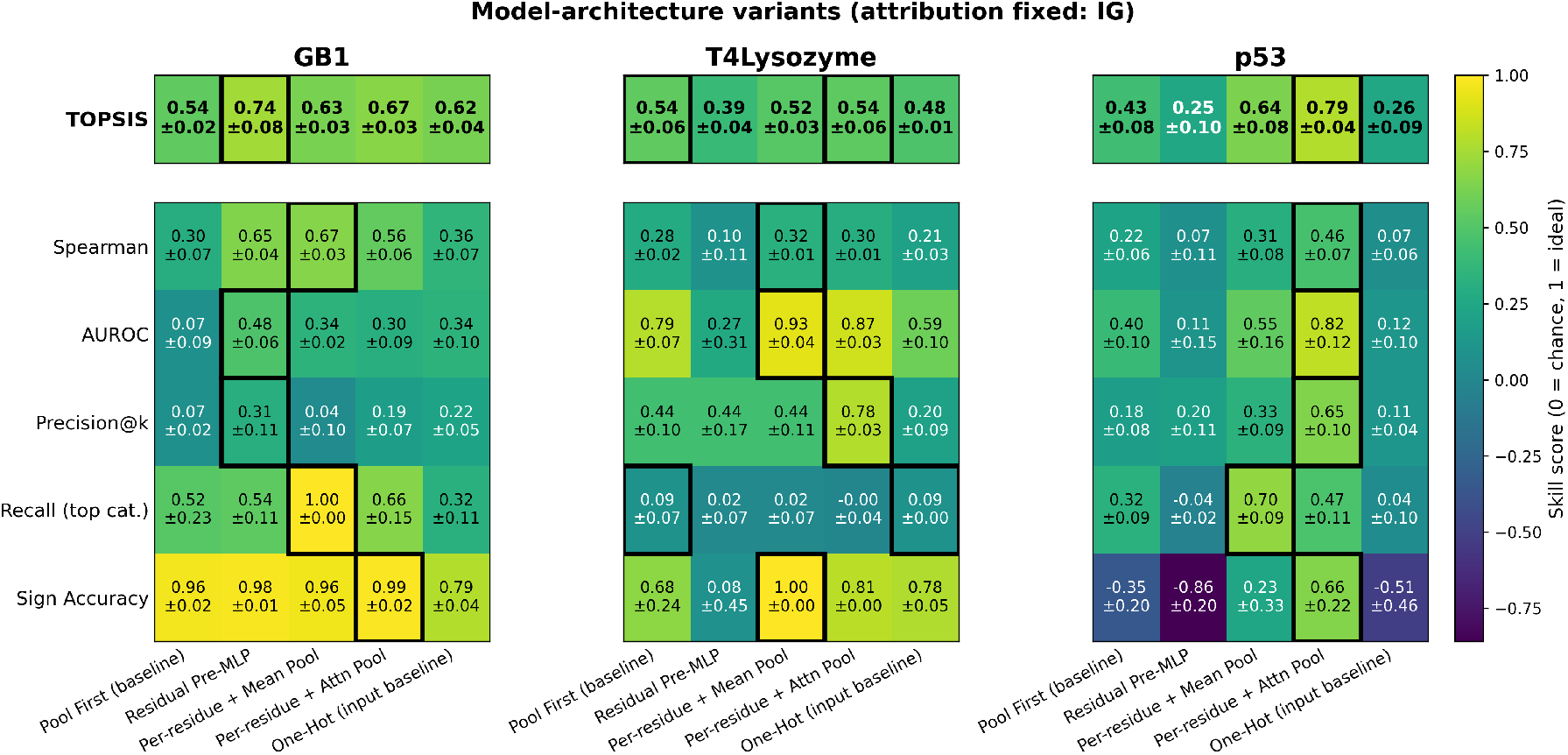
Performance breakdown of different architectures across synthetic proteins. Five architectures (Pool-First, Residual Pre-MLP, Mean Pool, Attn Pool, One-Hot control) on the ESM-C 300M backbone, scored per protein (GB1, T4 Lysozyme, p53) on five faithfulness metrics plus a composite TOPSIS score (top row). Values are mean ± SD across three synthetic landscapes, each at its best noise level, averaged over three common seeds. Color = skill score (0 = chance, 1 = ideal); black outlines mark the top-performing architecture(s) per row (ties included). Across all 27 protein landscape seed panels, a Friedman test confirms a significant overall difference in TOPSIS among architectures (*χ*^2^ = 16.65, *p* = 0.0023), and Attn Pool (mean TOPSIS = 0.665) significantly outperforms all four other architectures (Holm-corrected Wilcoxon, all *p <* 0.01).

**Fig. D.3.**
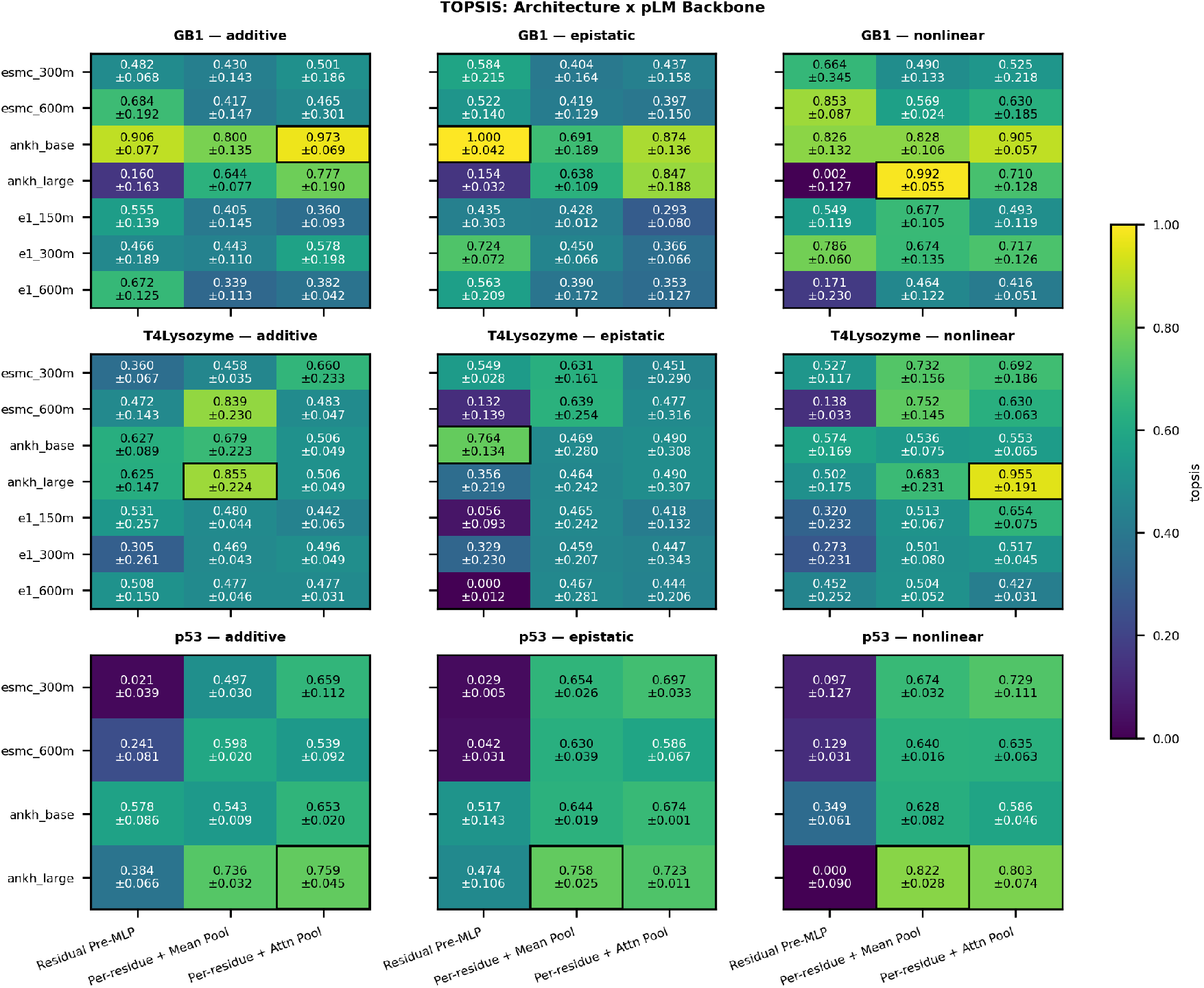
TOPSIS composite performance across architecture × pLM-backbone combinations. Each cell shows the mean ± seed-SD TOPSIS score at each combination’s best-performing noise level, aggregated across proteins and landscapes; the black outline marks the single best-performing combination overall. MeanPool+Ankh-large achieves the highest score (0.732, outlined), with AttnPool+Ankh-large essentially tied (0.730). A Friedman test confirms a significant overall difference among combinations (*χ*^2^ = 40.5, *p <* 0.0001), but pairwise Holm-corrected Wilcoxon tests (n=9 paired protein×landscape panels) only have power to detect the most extreme gap: MeanPool+Ankh-large significantly outperforms PreMLP+Ankh-large (*p* = 0.043). Other apparent gaps (e.g. vs. esmc-backbone or PreMLP combinations) do not survive multiple-comparison correction at this sample size and should not be read as equivalence.

**Fig. D.4.**
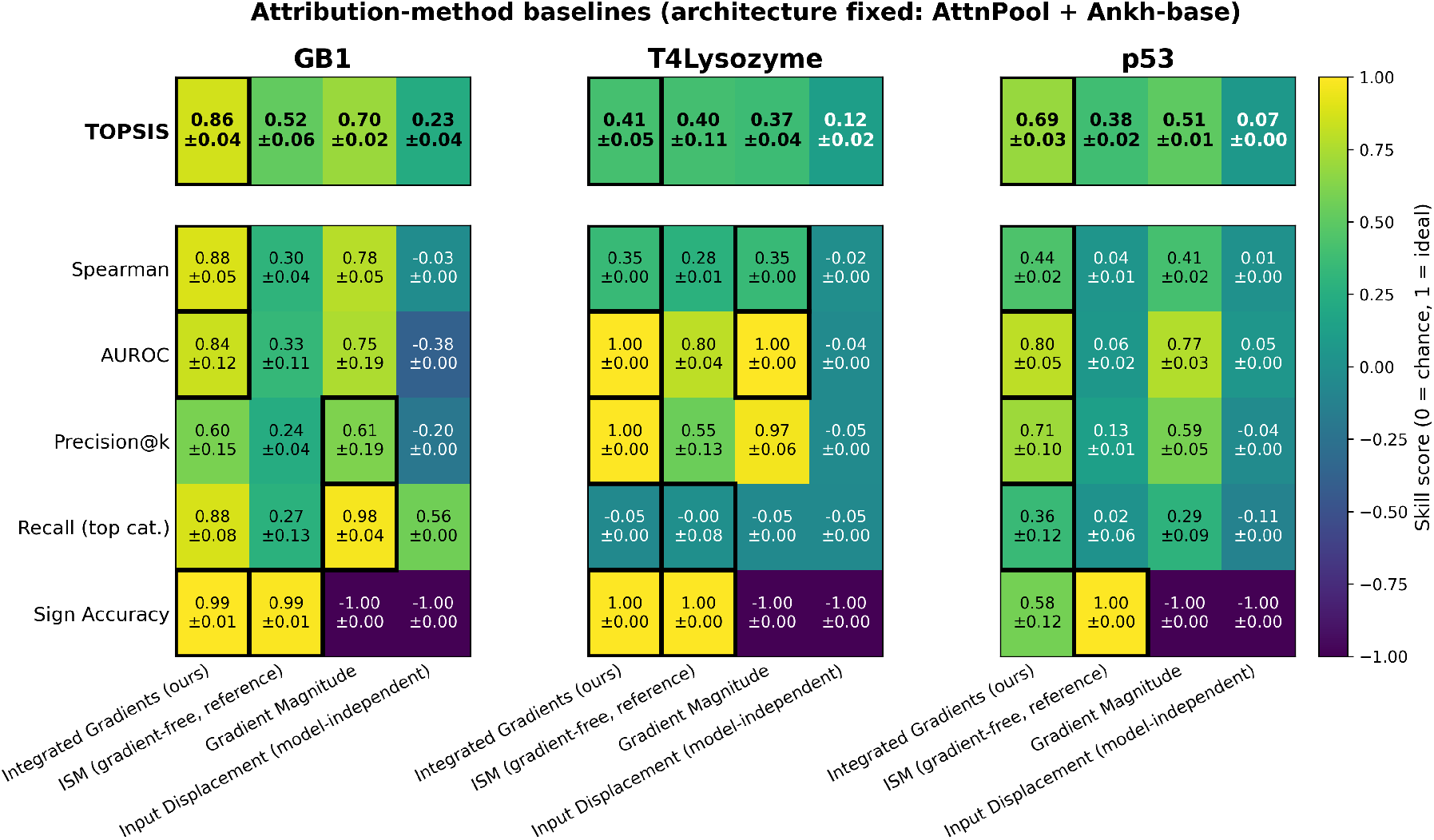
Metric breakdown for IG method and baselines. Comparison of Integrated Gradients (IG, ours) against ISM (gradient-free), Gradient Magnitude, and Input Displacement baselines, with architecture fixed to Per-residue + Attn Pool on Ankh-base embeddings. Each panel (GB1, T4 Lysozyme, p53) reports the five underlying skill metrics (Spearman, AUROC, Precision@*k*, Recall (top category), Sign Accuracy) and the composite TOPSIS score (top row), each at its own best noise level, averaged across the three synthetic landscapes (mean ± landscape-to-landscape SD). Color encodes skill score (0 = chance, 1 = ideal); black outlines mark the best-performing method(s) per row, with ties at the displayed precision included. Across all three proteins, IG achieves the highest mean TOPSIS score (0.65) and is significantly better than every baseline (Friedman *χ*^2^ = 74.3, *p <* 0.0001; Holm-corrected pairwise Wilcoxon, *p <* 0.0001 vs. each baseline).

## Appendix E Real Protein Results

## E.1 *α*-amylase

**Table E.1.** Attribution quality and model performance for the *α*-amylase splits (ESM per-residue mean pooling). Values are mean *±* s.d. over 5 seeds. Held-out Spearman *ρ*: correlation between predicted and measured activity on each split’s test set. Attribution metrics use the magnitude score 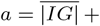 s.d.(*IG*) over all analyzed sequences against ground truth *w* = number of importance criteria met (Structural 2; Evolutionarily conserved and Detergent environment 1; Background 0); *k* = number of annotated positions (160 of 425); top category = Structural. ^†^Rescaled so that 0 is the mean for random attributions and 1 is perfect. ^‡^Shown as the skill score. Sign accuracy: fraction of annotated positions with mean IG *<* 0. TOPSIS: equal weights across the four splits. Recall rows: fraction of each category’s positions ranked in the top *k* (random = 0.38). Bold: best split for held-out *ρ* and TOPSIS.

|  | By mutation | One to many | Close to far | Far to close |
| --- | --- | --- | --- | --- |
| Held-out Spearman $\rho$ | **0.66 ± 0.01** | 0.65 ± 0.05 | 0.43 ± 0.02 | 0.38 ± 0.02 |
| TOPSIS | 0.77 | 0.43 | **0.91** | 0.00 |
| Spearman$^\dagger$ | 0.25 ± 0.03 | 0.16 ± 0.05 | 0.22 ± 0.02 | −0.01 ± 0.02 |
| AUROC$^\ddagger$ | 0.27 ± 0.03 | 0.17 ± 0.05 | 0.24 ± 0.02 | −0.01 ± 0.02 |
| Precision@$k^\dagger$ | 0.19 ± 0.03 | 0.13 ± 0.06 | 0.17 ± 0.01 | 0.01 ± 0.03 |
| Recall (top category)$^\dagger$ | 0.11 ± 0.08 | −0.02 ± 0.13 | 0.20 ± 0.03 | −0.03 ± 0.03 |
| Sign accuracy | 0.99 ± 0.01 | 0.99 ± 0.01 | 0.99 ± 0.01 | 1.00 ± 0.01 |
| *Recall in top k positions* |  |  |  |  |
| Structural + conserved | 0.45 ± 0.05 | 0.37 ± 0.08 | 0.50 ± 0.02 | 0.36 ± 0.02 |
| Conserved | 0.56 ± 0.01 | 0.53 ± 0.04 | 0.67 ± 0.01 | 0.23 ± 0.06 |
| Structural | 0.48 ± 0.02 | 0.46 ± 0.02 | 0.38 ± 0.01 | 0.46 ± 0.02 |
| Background | 0.31 ± 0.01 | 0.33 ± 0.02 | 0.31 ± 0.01 | 0.37 ± 0.01 |
| Sequences analyzed for IG | 3,707 | 3,707 | 3,707 | 3,707 |

**Fig. E.1.**
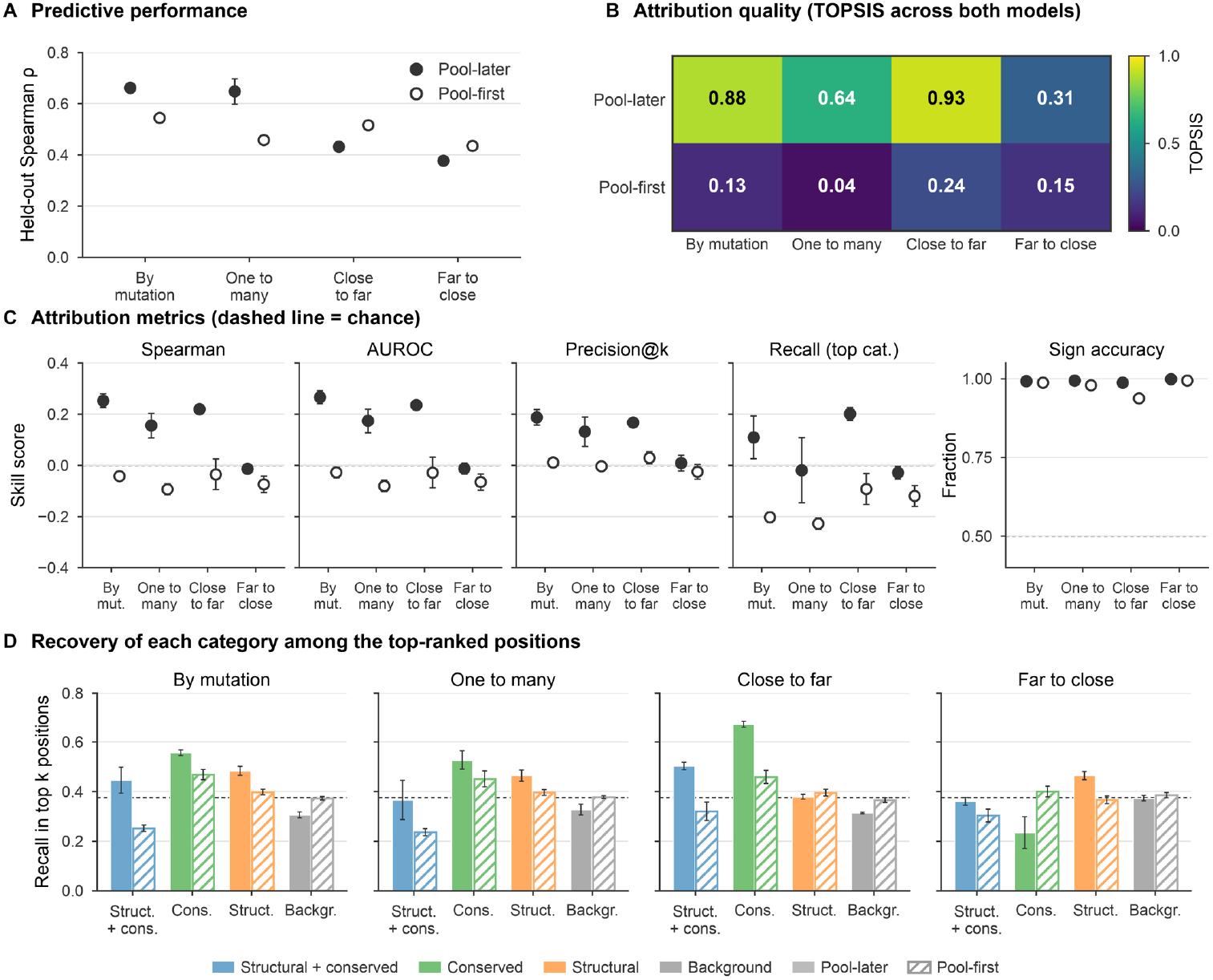
Pool-later models outperform pool-first *α*-amylase models in attribution quality. Pool-later (per-residue mean pooling; filled markers and bars) and pool-first (open markers, hatched bars) models on the four FLIP2 splits. Values are mean *±* s.d. over 5 seeds. (**A**) Held-out Spearman *ρ* between predicted and measured activity on each split’s test set. (**B**) TOPSIS score computed jointly over all eight model–split combinations with equal weights on the five attribution metrics in (C), so that scores are comparable across models. (**C**) Attribution metrics computed against the position categories defined in Section 4.9.4. Spearman, Precision@*k* and top-category recall are rescaled so that 0 is the mean for random attributions and 1 is perfect. Sign accuracy is the fraction of annotated positions with negative mean attribution (chance = 0.5). Dashed lines mark chance. (**D**) Fraction of each category’s positions ranked in the top *k* = 160 by attribution magnitude, where *k* is the number of annotated positions; the dashed line marks the random expectation (*k/N* = 0.38).

## E.2 *β*-lactamase

**Table E.2.** Rosetta ΔΔ*G*, Boltz-2 affinity, and integrated-gradient attributions for TEM-1 variants. ΔΔ*G*_Cart_ is the mean ± SD over 5 independent replicates (each with its own model FastRelax ensemble, lowest-scoring model selected); Boltz-2 values are the mean *±* SD over 20 independent runs.

| Dataset | Ligand | AA Substitution | Mean Integrated Gradients | Rosetta $\Delta\Delta G_{\text{Cart}}$ (kcal mol<sup>-1</sup>) | Boltz-2 Affinity Pred log<sub>10</sub> IC<sub>50</sub> (μM) |
| --- | --- | --- | --- | --- | --- |
| Deng 2012 | ampicillin | A235K | -2.602 | 0.29 ± 0.01 | -0.07 ± 0.06 |
| Deng 2012 | ampicillin | A235R | -2.548 | 0.35 ± 0.01 | 0.03 ± 0.07 |
| Deng 2012 | ampicillin | M115W | -4.426 | 4.86 ± 0.21 | -0.80 ± 0.07 |
| Deng 2012 | ampicillin | E210K | 0.087 | -0.27 ± 0.03 | -0.79 ± 0.04 |
| Deng 2012 | ampicillin | E210I | 0.087 | -0.34 ± 0.10 | -0.79 ± 0.08 |
| Deng 2012 | ampicillin | E210R | 0.086 | -0.45 ± 0.09 | -0.76 ± 0.08 |
| Deng 2012 | ampicillin | T116F | -2.878 | 1.49 ± 0.04 | -0.70 ± 0.09 |
| Deng 2012 | ampicillin | T116C | -2.862 | 1.41 ± 0.09 | -0.70 ± 0.10 |
| Deng 2012 | ampicillin | D83V | -3.250 | 1.64 ± 0.05 | -0.73 ± 0.10 |
| Deng 2012 | ampicillin | D83P | -3.238 | 9.80 ± 0.16 | -0.76 ± 0.05 |
| Deng 2012 | ampicillin | D83R | -3.230 | 2.53 ± 0.21 | -0.75 ± 0.09 |
| Firnberg 2014 | ampicillin | D129H | -1.105 | 8.81 ± 0.57 | -0.54 ± 0.08 |
| Firnberg 2014 | ampicillin | D129R | -1.093 | 19.58 ± 0.24 | -0.42 ± 0.08 |
| Firnberg 2014 | ampicillin | D129G | -1.090 | 6.25 ± 0.05 | -0.60 ± 0.11 |
| Firnberg 2014 | ampicillin | V78W | -1.123 | 7.73 ± 0.24 | -0.75 ± 0.06 |
| Firnberg 2014 | ampicillin | E56P | -0.758 | 8.32 ± 0.17 | -0.76 ± 0.06 |
| Firnberg 2014 | ampicillin | V29M | 0.001 | 0.18 ± 0.21 | -0.73 ± 0.07 |
| Firnberg 2014 | ampicillin | V31L | 0.005 | 2.60 ± 0.42 | -0.73 ± 0.07 |
| Firnberg 2014 | ampicillin | V31I | 0.006 | 0.91 ± 0.24 | -0.74 ± 0.10 |
| Jacquier 2013 | amoxicillin | V257E | -1.393 | 4.57 ± 0.50 | -0.71 ± 0.04 |
| Jacquier 2013 | amoxicillin | D212N | 0.186 | 1.20 ± 0.60 | -0.72 ± 0.03 |
| Jacquier 2013 | amoxicillin | D212V | 0.107 | 0.22 ± 0.13 | -0.69 ± 0.03 |
| Jacquier 2013 | amoxicillin | R238G | -0.304 | 0.51 ± 0.02 | -0.67 ± 0.04 |
| Jacquier 2013 | amoxicillin | V29E | 0.113 | 0.23 ± 0.20 | -0.71 ± 0.03 |
| Jacquier 2013 | amoxicillin | M180T | 0.188 | 0.11 ± 0.06 | -0.77 ± 0.04 |
| Stiffler 2015 | ampicillin | K71Y | -3.152 | -0.50 ± 0.34 | -0.83 ± 0.05 |
| Stiffler 2015 | cefotaxime | K71Y | -3.152 | -0.50 ± 0.34 | -1.29 ± 0.05 |
| Stiffler 2015 | ampicillin | C75H | -2.864 | 2.51 ± 0.12 | -0.78 ± 0.08 |
| Stiffler 2015 | cefotaxime | C75H | -2.864 | 2.51 ± 0.12 | -1.30 ± 0.05 |
| Stiffler 2015 | ampicillin | V72F | -2.613 | 3.78 ± 0.70 | -0.85 ± 0.07 |
| Stiffler 2015 | cefotaxime | V72F | -2.613 | 3.78 ± 0.70 | -1.34 ± 0.04 |
| Stiffler 2015 | ampicillin | A235S | 0.008 | 0.08 ± 0.01 | -0.32 ± 0.11 |
| Stiffler 2015 | cefotaxime | A235S | 0.008 | 0.08 ± 0.01 | -1.29 ± 0.05 |
| Stiffler 2015 | ampicillin | F64Y | -1.502 | 0.59 ± 0.02 | -0.72 ± 0.09 |
| Stiffler 2015 | cefotaxime | F64Y | -1.502 | 0.59 ± 0.02 | -1.26 ± 0.11 |

**Table E.3.** Functional annotations of tested residue positions in TEM-1 *β*-lactamase (BLAT).

| Ambler | ProteinGym | WT AA | Annotation | Functional Annotation | Source |
| --- | --- | --- | --- | --- | --- |
| 31 | 29 | Val (V) | Structural | Substitution increases stability | Palzkill 2018 |
| 66 | 64 | Phe (F) | Structural | Beta-bridge packing against T180–M182 | Philippon et al. 2016 |
| 73 | 71 | Lys (K) | Catalytic | General base of the S-x-x-K motif, deprotonating S70 for acylation | Palzkill 2018 |
| 77 | 75 | Cys (C) | Disulfide | Forms the single disulfide bond of TEM-1 with C123 | UniProt P62593 |
| 81 | 79 | Leu (L) | Structural | 95% conserved buried hydrophobic core packing in helix H2 | Philippon et al. 2016 |
| 131 | 129 | Asp (D) | Structural | Fully conserved helix H5 N-cap that holds SDN-loop geometry | Philippon et al. 2016 |
| 182 | 180 | Met (M) | Structural | Global-stability hot spot whose M182T substitution raises $T_m$ by 7 °C. | Palzkill 2018 |
| 214 | 212 | Asp (D) | Structural | Second-shell hydrogen-bond partner of D233 in loop 213–220 | Philippon et al. 2016 |
| 237 | 235 | Ala (A) | Catalytic | Main-chain amide forms the oxyanion hole with that of catalytic S70 | Palzkill 2018 |
| 241 | 238 | Arg (R) | Binding | R241G slightly improves catalytic efficiency | Venkatachalam et al. 1994 |

## References

[1] Notin, P., Rollins, N., Gal, Y., Sander, C., Marks, D.: Machine learning for functional protein design. Nature Biotechnology 42(2), 216–228 (2024) 10.1038/s41587-024-02127-0 . Accessed 2026-10-21

[2] Barghout, R.A., Kirby, M., Zheng, A., Chinas, L., Mohammadi, M., Xu, Z., Sanchez-Lengeling, B., Mahadevan, R.: Machine Learning for Sequence-to-Function Approaches. In: Machine Learning and Big Data-enabled Biotechnology, pp. 103–129. John Wiley & Sons, Ltd, Online (2026). 10.1002/9783527850532.ch4

[3] Notin, P., Kollasch, A., Ritter, D., Niekerk, L., Paul, S., Spinner, H., Rollins, N., Shaw, A., Orenbuch, R., Weitzman, R., Frazer, J., Dias, M., Franceschi, D., Gal, Y., Marks, D.: ProteinGym: Large-Scale Benchmarks for Protein Fitness Prediction and Design. In: Oh, A., Naumann, T., Globerson, A., Saenko, K., Hardt, M., Levine, S. (eds.) Advances in Neural Information Processing Systems, vol. 36, pp. 64331–64379 (2023). 10.52202/075280-2810

[4] Dallago, C., Mou, J., Johnston, K.E., Wittmann, B.J., Bhattacharya, N., Goldman, S., Madani, A., Yang, K.K.: FLIP: Benchmark tasks in fitness landscape inference for proteins. In: Vanschoren, J., Yeung, S. (eds.) Proceedings of the Neural Information Processing Systems Track on Datasets and Benchmarks, vol. 1 (2021). https://datasets-benchmarks-proceedings.neurips.cc/paper_files/paper/2021/file/2b44928ae11fb9384c4cf38708677c48-Paper-round2.pdf

[5] Didi, K., Alamdari, S., Lu, A.X., Wittmann, B., Johnston, K.E., Amini, A.A., Madani, A., Czeneszew, M., Dallago, C., Yang, K.K.: Flip2: Expanding protein fitness landscape benchmarks for real-world machine learning applications. bioRxiv (2026) 10.64898/2026.02.23.707496

[6] Sundararajan, M., Taly, A., Yan, Q.: Axiomatic Attribution for Deep Networks. In: Proceedings of the 34th International Conference on Machine Learning. Proceedings of Machine Learning Research, vol. 70, pp. 3319–3328 (2017). https://proceedings.mlr.press/v70/sundararajan17a.html

[7] Sturmfels, P., Lundberg, S., Lee, S.-I.: Visualizing the impact of feature attribution baselines. Distill (2020) 10.23915/distill.00022 . https://distill.pub/2020/attribution-baselines

[8] Zaher, E., Trzaskowski, M., Nguyen, Q., Roosta, F.: Manifold Integrated Gradients: Riemannian Geometry for Feature Attribution. In: Proceedings of the 41st International Conference on Machine Learning. Proceedings of Machine Learning Research, vol. 235, pp. 58090–58104 (2024). https://proceedings.mlr.press/v235/zaher24a.html

[9] Miglani, V., Kokhlikyan, N., Alsallakh, B., Martin, M., Reblitz-Richardson, O.: Investigating Saturation Effects in Integrated Gradients. arXiv. arXiv:2010.12697 [cs]. Presented at the ICML Workshop on Human Interpretability in Machine Learning (WHI 2020) (2020). 10.48550/arXiv.2010.12697

[10] Walker, C., Jha, S., Chen, K., Ewetz, R.: Integrated Decision Gradients: Compute Your Attributions Where the Model Makes Its Decision. AAAI 38(6), 5289–5297 (2024) 10.1609/aaai.v38i6.28336

[11] Agarwal, V., Shendure, J.: Predicting mRNA Abundance Directly from Genomic Sequence Using Deep Convolutional Neural Networks. Cell Reports 31(7), 107663 (2020) 10.1016/j.celrep.2020.107663

[12] Jha, A., Aicher, J.K., Gazzara, M.R., Singh, D., Barash, Y.: Enhanced Integrated Gradients: improving interpretability of deep learning models using splicing codes as a case study. Genome Biol 21(1), 149 (2020) 10.1186/s13059-020-02055-7

[13] Valencia, J.D., Hendrix, D.A.: Improving deep models of protein-coding potential with a Fourier-transform architecture and machine translation task. PLOS Computational Biology 19(10), 1011526 (2023) 10.1371/journal.pcbi.1011526

[14] Hunklinger, A., Ferruz, N.: Toward the Explainability of Protein Language Models. arXiv. arXiv:2506.19532 [q-bio] (2025). 10.48550/arXiv.2506.19532

[15] Simon, E., Zou, J.: InterPLM: discovering interpretable features in protein language models via sparse autoencoders. Nature Methods 22(10), 2107–2117 (2025) 10.1038/s41592-025-02836-7

[16] Fazel, Z., De Souza, C.P.E., Golding, G.B., Ilie, L.: Explainability of Protein Deep Learning Models. IJMS 26(11), 5255 (2025) 10.3390/ijms26115255

[17] Wenzel, M., Grüner, E., Strodthoff, N.: Insights into the inner workings of transformer models for protein function prediction. Bioinformatics 40(3), 031 (2024) 10.1093/bioinformatics/btae031

[18] Del Giudice, M.: The prediction-explanation fallacy: A pervasive problem in scientific applications of machine learning. Methodology: European Journal of Research Methods for the Behavioral and Social Sciences 20(1), 22–46 (2024) 10.5964/meth.11235

[19] Yao, J., Song, A., Baerenfaller, K., Zhakparov, D.: Residue-Level Attributions in Protein Language Models Do Not Recover Allergen Epitopes. arXiv. arXiv:2606.22181 [cs.LG]. Accepted at the ICML 2026 Mechanistic Interpretability Workshop (2026). 10.48550/arXiv.2606.22181

[20] Samek, W., Binder, A., Montavon, G., Lapuschkin, S., Müller, K.-R.: Evaluating the visualization of what a Deep Neural Network has learned. IEEE Transactions on Neural Networks and Learning Systems 28(11), 2660–2673 (2017) 10.1109/TNNLS.2016.2599820

[21] Hooker, S., Erhan, D., Kindermans, P.-J., Kim, B.: A Benchmark for Interpretability Methods in Deep Neural Networks. In: Wallach, H., Larochelle, H., Beygelzimer, A., d’Alché-Buc, F., Fox, E., Garnett, R. (eds.) Advances in Neural Information Processing Systems, vol. 32 (2019). https://proceedings.neurips.cc/paper_files/paper/2019/hash/fe4b8556000d0f0cae99daa5c5c5a410-Abstract.html

[22] DeYoung, J., Jain, S., Rajani, N.F., Lehman, E., Xiong, C., Socher, R., Wallace, B.C.: ERASER: A Benchmark to Evaluate Rationalized NLP Models. In: Jurafsky, D., Chai, J., Schluter, N., Tetreault, J. (eds.) Proceedings of the 58th Annual Meeting of the Association for Computational Linguistics, pp. 4443–4458. Association for Computational Linguistics, Online (2020). 10.18653/v1/2020.acl-main.408

[23] Barraza-Chavez, J.M., Barghout, R.A., Almada-Monter, R., Sanchez-Lengeling, B., Jinich, A., Mahadevan, R.: Graph Data Modeling: Molecules, Proteins, & Chemical Processes. American Chemical Society, Online (2025). 10.1021/acsinfocus.7e9017 . https://pubs.acs.org/infocus/monograph/4288/Graph-Data-Modeling-Molecules-Proteins-amp Accessed 2026-10-20

[24] Sanchez-Lengeling, B., Wei, J., Lee, B., Reif, E., Wang, P., Qian, W., McCloskey, K., Colwell, L., Wiltschko, A.: Evaluating Attribution for Graph Neural Networks. In: Advances in Neural Information Processing Systems, vol. 33, pp. 5898–5910. Curran Associates, Inc., Online (2020). https://proceedings.neurips.cc/paper/2020/hash/417fbbf2e9d5a28a855a11894b2e795a-Abstract.html Accessed 2026-10-20

[25] Chernigovskaya, M., Pavlović, M., Kanduri, C., Gielis, S., Robert, P., Scheffer, L., Slabodkin, A., Haff, I.H., Meysman, P., Yaari, G., Sandve, G.K., Greiff, V.: Simulation of adaptive immune receptors and repertoires with complex immune information to guide the development and benchmarking of AIRR machine learning. Nucleic Acids Research 53(3), 025 (2025) 10.1093/nar/gkaf025 . Accessed 2026-10-21

[26] ESM Team: ESM Cambrian: Revealing the mysteries of proteins with unsupervised learning. EvolutionaryScale Website (2024). https://evolutionaryscale.ai/blog/esm-cambrian

[27] Jain, S., Beazer, J., Ruffolo, J.A., Bhatnagar, A., Madani, A.: E1: Retrieval-Augmented Protein Encoder Models. bioRxiv (2025). 10.1101/2025.11.12.688125

[28] Elnaggar, A., Essam, H., Salah-Eldin, W., Moustafa, W., Elkerdawy, M., Rochereau, C., Rost, B.: Ankh: Optimized Protein Language Model Unlocks General-Purpose Modelling. arXiv. arXiv:2301.06568 [cs] (2023). 10.48550/arXiv.2301.06568

[29] Alsamkary, H., Elshaffei, M., Elkerdawy, M., Elnaggar, A.: Ankh3: Multi-Task Pretraining with Sequence Denoising and Completion Enhances Protein Representations. arXiv (2025). 10.48550/arXiv.2505.20052

[30] Yu, T., Cui, H., Li, J.C., Luo, Y., Jiang, G., Zhao, H.: Enzyme function prediction using contrastive learning. Science 379(6639), 1358–1363 (2023) 10.1126/science.adf2465

[31] Yu, H., Deng, H., He, J., Keasling, J.D., Luo, X.: UniKP: a unified framework for the prediction of enzyme kinetic parameters. Nature Communications 14(1), 8211 (2023) 10.1038/s41467-023-44113-1

[32] Adebayo, J., Gilmer, J., Muelly, M., Goodfellow, I., Hardt, M., Kim, B.: Sanity Checks for Saliency Maps. In: Bengio, S., Wallach, H., Larochelle, H., Grauman, K., Cesa-Bianchi, N., Garnett, R. (eds.) Advances in Neural Information Processing Systems, vol. 31 (2018). https://proceedings.neurips.cc/paper_files/paper/2018/hash/294a8ed24b1ad22ec2e7efea049b8737-Abstract.html

[33] Li, F.-Z., Amini, A.P., Yue, Y., Yang, K.K., Lu, A.X.: Feature Reuse and Scaling: Understanding Transfer Learning with Protein Language Models. bioRxiv (2024). 10.1101/2024.02.05.578959

[34] West-Roberts, J., Kravitz, J., Jha, N., Cornman, A., Hwang, Y.: Diverse Genomic Embedding Benchmark for functional evaluation across the tree of life. bioRxiv (2024). 10.1101/2024.07.10.602933

[35] Kumar, A., Jha, I.: Layer Probing Improves Kinase Functional Prediction with Protein Language Models. arXiv. arXiv:2512.00376 [q-bio.QM] (2025). 10.48550/arXiv.2512.00376

[36] Hart, A., Han, C., Kim, J., Zhao, H., Ji, H.: Protein Language Models Diverge from Natural Language: Comparative Analysis and Improved Inference. arXiv. arXiv:2602.20449 [cs.LG] (2026). 10.48550/arXiv.2602.20449

[37] Passaro, S., Corso, G., Wohlwend, J., Reveiz, M., Thaler, S., Somnath, V.R., Getz, N., Portnoi, T., Roy, J., Stark, H., Kwabi-Addo, D., Beaini, D., Jaakkola, T., Barzilay, R.: Boltz-2: Towards Accurate and Efficient Binding Affinity Prediction. bioRxiv (2025). 10.1101/2025.06.14.659707

[38] Hwang, C.-L., Yoon, K.: Multiple Attribute Decision Making: Methods and Applications. Lecture Notes in Economics and Mathematical Systems, vol. 186. Springer, Berlin, Heidelberg (1981). 10.1007/978-3-642-48318-9

[39] Deng, Z., Huang, W., Bakkalbasi, E., Brown, N.G., Adamski, C.J., Rice, K., Muzny, D., Gibbs, R.A., Palzkill, T.: Deep sequencing of systematic combinatorial libraries reveals β-lactamase sequence constraints at high resolution. Journal of Molecular Biology 424(3-4), 150–167 (2012) 10.1016/j.jmb.2012.09.014

[40] Firnberg, E., Labonte, J.W., Gray, J.J., Ostermeier, M.: A comprehensive, high-resolution map of a gene’s fitness landscape. Molecular Biology and Evolution 31(6), 1581–1592 (2014) 10.1093/molbev/msu081

[41] Jacquier, H., Birgy, A., Le Nagard, H., Mechulam, Y., Schmitt, E., Glodt, J., Bercot, B., Petit, E., Poulain, J., Barnaud, G., Gros, P.-A., Tenaillon, O.: Capturing the mutational landscape of the beta-lactamase TEM-1. Proceedings of the National Academy of Sciences of the United States of America 110(32), 13067–13072 (2013) 10.1073/pnas.1215206110

[42] Stiffler, M.A., Hekstra, D.R., Ranganathan, R.: Evolvability as a function of purifying selection in TEM-1 β-lactamase. Cell 160(5), 882–892 (2015) 10.1016/j.cell.2015.01.035

